# Multiomic evidence for microbial contributions to species-specific olfactory communication in wild tamarins

**DOI:** 10.64898/2026.10.02.756344

**Authors:** Silvia Carboni, Alice C. Poirier, Mrinalini Watsa, Gideon A. Erkenswick, Amanda D. Melin

**Affiliations:** Department of Anthropology & Archaeology, University of Calgary, 2500 University Dr. NW, Calgary, AB T2N 1N4, Canada; Université Bourgogne Europe, Institut Agro, CNRS, INRAE, UMR CSGA, Development of Olfactory Communication and Cognition Laboratory, Dijon, 21000, France; Conservation Science and Wildlife Health, San Diego Zoo Wildlife Alliance, Escondido, California, USA; Field Projects International, Escondido, California, USA; Department of Medical Genetics, University of Calgary, 2500 University Dr NW, Calgary, AB T2N 1N4, Canada; Alberta Children’s Hospital Research Institute, University of Calgary, 3330 Hospital Dr, NW, Calgary, AB T2N 4N1, Canada

**Keywords:** scent gland microbiome, volatile organic compounds, reproductive isolation, social communication

## Abstract

Closely related sympatric species are hypothesized to face strong selection against hybridization, favoring species-specific signals that maintain reproductive isolation. Within species, olfaction may also mediate social interactions and facilitate social cohesion. Here, we investigated the potential for microbiome-derived scent signals to contribute to inter- and intra-species communication in two sympatric wild tamarins (*Leontocebus weddelli* and *Tamarinus subgrisescens*). We analyzed scent gland samples to characterize odor profiles (volatile organic compounds, VOCs) and microbiome taxonomic composition and putative functions from metagenomic shotgun sequences. Species membership was predicted accurately from microbiome profiles, whereas VOCs alone discriminated poorly between species. We found high abundance of *Corynebacterium* in *L. weddelli*, a genus associated with fatty acid metabolism, alongside the exclusive detection of acetic acid, highlighting microbially derived fatty acids as candidates for further investigation in this species. In *T. subgrisescens,* we found a more diverse microbiota, with fewer taxa known to be related to odor production, while VOC profiles appeared less variable among individuals. In contrast, group-level signatures were weak, suggesting that intra-specific odor may not strongly contribute to signaling group membership. We highlight the challenge of linking high-dimensional microbial communities, which include taxa not necessarily involved in odor production, to the smaller dataset of detectable VOCs. Although scent gland microbiota differed between tamarin species, whether these differences translate into species-specific odor cues remains to be tested. Future research involving behavioural assays and targeted experimental approaches that focus on key microbes and metabolites may shed further insight into roles and mechanisms of olfactory communication.

## INTRODUCTION

Understanding the processes leading to reproductive isolation and, ultimately, speciation, has been a longstanding goal in evolutionary biology (Janicke et al., 2019; Ptacek, 2000). Selection against hybrids has been proposed as one of the driving mechanisms, favoring traits that reduce the production of offspring with intermediate phenotypes maladapted to parental ecological niches (Thompson et al., 2024). Assortative mating (i.e., preference for more phenotypically similar partners) has been proposed as a mechanism of speciation, especially in sympatric populations where spatial barriers are not a constraint for gene flow and hybridization (Janicke et al., 2019; Lawson et al., 2023; Muralidhar et al., 2022; Winters & Higham, 2022). Despite their relevance for advancing the field, the proximate mechanisms through which populations maintain reproductive isolation remain poorly known. Nevertheless, sensory systems are considered central to these mechanisms.

Sensory systems define how organisms perceive and interact with the world around them, playing a key role in identifying conspecifics and allospecifics. Studies of inter-species phenotypic diversity in sensory traits often complement research efforts to understand speciation mechanisms (Garcia et al., 2020; Ptacek, 2000; Van Huynh & Rice, 2025; Wilkins et al., 2013; Winters & Higham, 2022). For highly visual species such as primates, research in this area has mainly focused on colourful traits and character displacement perceived through visual communication (Allen et al., 2014; Laurita & Poindexter, 2025; Santana et al., 2012), but some studies have also explored the acoustic potential for species differentiation (Braune et al., 2008; Hasiniaina et al., 2020; Markolf Rakotonirina, 2017). Growing evidence suggests that olfactory cues may also contribute to species recognition (Coombes et al., 2018; Epple et al., 1987; Lang et al., 2025; K. C. Rowe et al., 2020; Van Huynh & Rice, 2025), despite having historically been overlooked as a key sensory system, particularly in monkeys, apes, and humans (haplorhine primates). Yet, the role of olfaction in haplorhine primates is increasingly recognized (e.g., Henkel & Setchell, 2018; Heymann, 2022; Poirier & Melin, 2024; Simon et al., n.d.). As with visual and acoustic cues, olfaction may facilitate species discrimination and contribute to assortative mating, potentially reinforcing reproductive isolation in species that share territorial boundaries.

In addition to species-level recognition, communication of group membership to conspecifics is key in social animals (Ward & Webster, 2016). Inter-group interactions range from peaceful to highly aggressive in mammals, but even among relatively non-aggressive interactions, communicating group membership may be relevant for mediating access to shared resources, such as mating opportunities, territory, and food (De Dreu & Triki, 2022; Ward & Webster, 2016). Olfactory communication may contribute to regulating these interactions by facilitating group member recognition (Brennan & Kendrick, 2006). For example, familiarity with group-specific odors may promote social affiliation, agonism, and influence mate preferences (Coombes et al., 2018; Fisher et al., 2003; Schubert et al., 2026). Yet, the role of olfactory cues in mediating social communication within and between groups remains poorly understood.

Studies of mammalian olfactory communication and body odor production have historically been hindered by technical limitations. However, recent advances in sequencing and molecular technologies now enable more robust tests of evolutionary hypotheses of body odor production. One of the most prominent, the fermentation hypothesis, proposes that symbiotic microbes inhabiting mammalian skin scent glands significantly contribute to body odor by releasing volatile organic compounds (VOCs) as by-products of their metabolism (Albone & Shirley, 1984; Carthey et al., 2018). In hosts where olfaction communicates reproductive isolation and group membership, natural selection may favor the development and maintenance of species-specific and group-specific glandular microbiomes and associated VOCs as an adaptive trait (Shropshire & Bordenstein, 2016). Past studies have attempted to disentangle the complex interactions between hosts, microbiome, and odors by testing whether bacterial taxonomic diversity positively associates with VOC diversity across vertebrates, including lemurs (Greene et al., 2019), carnivores (Leclaire et al., 2017; Rojas et al., 2023; Theis et al., 2013), pandas (Ma et al., 2022), and birds (Whittaker et al., 2019). These studies highlight two main gaps. First, they largely focus on the bacterial component of the microbiota, overlooking other taxa that may contribute to odor profiles (e.g., eukaryotes). Second, studies of the functional potential of microbial communities remain less common, despite potentially providing a more direct link between microbiomes and VOC production. While taxonomic analyses identify the microorganisms, functional pathway analyses can reveal the metabolic processes responsible for producing odor compounds.

In the present research, we capitalize on advances in sequencing technologies that enable comprehensive community profiling and functional characterization. We combine multiomic datasets to assess the putative influence of glandular microbiome on VOC production, ultimately aiming to elucidate the role of olfaction in communicating species and group identity. We study two sympatric species of wild tamarin monkeys (*Leontocebus weddelli* and *Tamarinus subgrisescens*), highly social primates in the Amazon basin. Tamarins present specialized histological and behavioral traits linked to olfactory communication: three discrete scent glands and scent marking (i.e., the act of depositing odor cues in the environment) (Epple et al., 1986; Heymann, 2006; Moraes et al., 2006). Tamarin scent glands harbor a microbiome capable of contributing to odor production (Carboni et al., 2025), and the maintenance of these energetically costly traits suggests that olfaction plays a role in social and reproductive communication (Carboni et al., 2026a; Heymann, 1998, 2022). Our hypotheses are that scent gland microbiomes and VOCs are involved in: 1) mediating inter-specific discrimination, potentially contributing to reproductive isolation, and 2) communicating group membership, potentially contributing to intra-specific social cohesion. We predicted that i) covariance among scent gland microbial taxa and VOCs, and among microbial metabolic pathways and VOCs, will exhibit a species-specific structure, and that ii) microbiome and VOC diversity and composition will differ between social groups of the same species. To test our predictions, we generated three connected datasets for multiple social groups of each species: scent gland microbial taxonomy based on shotgun metagenomic sequencing, microbial metabolic pathways predicted from gene families identified from the metagenomic data, and VOC profiles from thermal desorption–gas chromatography-mass spectrometry.

## METHODS

### Study Site and Study Species

This study was conducted in the Madre de Dios Department of southeast Peru, at the Estación Biológica Río Los Amigos (EBLA). The research station is located within a conservation area managed by Conservación Amazónica-ACCA, which experiences a wet season (October to March) and a dry season (April to September), with stable average temperatures ranging from 21°C to 25°C (Watsa, 2013). The area is inhabited by our two study species: Weddell’s saddleback tamarins (*Leontocebus weddelli*) and bearded emperor tamarins (*Tamarinus subgrisescens*). The tamarin population has been monitored by the research organization Field Projects International (FPI) since 2009, providing long-term demographic data. Both species are small (*L. weddelli*: 394 ± 31g; *T. subgrisescens*: 531 ± 67g), polyandrous, living in social groups of 3 to 9 individuals, typically consisting of a breeding female, young offspring, multiple males and occasionally non-breeding females (Watsa, 2013).

The two focal tamarin species are well suited for our study for two reasons. First, they overlap extensively in habitat use and are frequently observed travelling and foraging in close proximity (Watsa, 2013; Windfelder, 1997), conditions that could facilitate hybridization. Although no cases of inter-specific mating have been reported in this population, they share a recent evolutionary history (Brcko et al., 2022), and hybridization has been reported in several mammals under similar conditions (Adavoudi & Pilot, 2021; Detwiler et al., 2005; Neaves et al., 2010), including in primates from different genera (Zinner et al., 2011), and in tamarins (Pérez-Estrada & Soto-Calderón, 2026; Soto-Calderón et al., 2018). Given that hybrids are often associated with reduced fitness (Adavoudi & Pilot, 2021; Thompson et al., 2024), mechanisms involved in the maintenance of species boundaries may have evolved in this system. Second, both species are cooperative breeders, and their reproductive success depends heavily on alloparental care (Bales et al., 2000). Such a system may increase the fitness benefits of sharing resources with familiar group members while reducing investment in other individuals (Bergmüller et al., 2007; Diaz-Munoz, 2011). Under these conditions, selection may favor mechanisms that enable individuals to discriminate among group members and conspecific outsiders.

### Sample collection

We collected samples between June and July in 2022, 2023, and 2024 from 7 groups of *L. weddelli* and 7 groups of *T. subgrisescens* (Suppl. Materials 1), capitalizing on the annual mark-recapture program led by FPI, whose protocols are publicly available (Watsa et al., 2023). Individuals are longitudinally identified through subcutaneous microchips, and we collected non-invasive microbial and volatile organic compound (VOC) samples from the suprapubic glands of each individual trapped at most once per year. We first collected glandular VOCs using dynamic headspace extraction into custom-made glass probes filled with a mix of absorbent polymers (1.5 mg Tenax, 1.5 mg Carbotrap, 1.5 mg Carbosieve III) connected by silicone tubing to a portable air pump (Nevo & Valenta, 2018). To capture the VOCs, we pumped the air above the gland, ∼1 cm above the skin, then promptly placed the probe into an individual vial (2 mL with 0.3 mL fused insert) sealed with a Teflon screw cap. In 2022, the air pump used was custom-made, set to its minimum flow rate of 1.5 L/min, and sampled for 4 min. In 2023-24, we replaced it with an SKC Pocket Pump^®^ Touch, set to a slower flow rate and a longer extraction (0.5 L/min for 5 min) to facilitate VOC retention into the glass probes.

Immediately after VOCs collection, we swabbed the glandular surface area for 10 seconds using dry swabs (BD BBL CultureSwabTM) as previously described in Carboni et al. (2025). For each trapping day, we also collected one swab and two probes as ambient field controls, following the same protocol and sampling the surrounding air instead of the animal. Vials and swabs were placed on ice immediately after collection and stored in the camp freezer (-25°C) within 6 hours of collection.

### Microbiome analysis

We extracted DNA from all swab samples within a month of collection at the BSL-2 genetic lab located on site (Los Amigos Wildlife Conservation Laboratory - In Situ Lab Initiative). We used the DNeasy® PowerSoil Pro Kit with spin columns (Qiagen ID. 47017) following the manufacturer’s protocol except for skipping the steps with CD2 buffer to prevent DNA loss in our low-concentration samples. At each extraction event, we included laboratory controls following best standards in low-biomass microbial studies (Eisenhofer et al., 2019; Kim et al., 2017). Samples were stored at -25°C until shipment to the University of Calgary. Libraries were prepared using the NEB Ultra II DNA library prep kit; samples collected in 2022 were processed at the Center for Health Genomics and Informatics, while the libraries of samples from subsequent years were prepared at the Melin Genomic Lab. We included 6 field negative controls, 24 laboratory negative controls, and one positive control mock community from the ZymoBIOMICS Microbial Community Standard (D6300, Zymo-Research, Irvine, CA, USA). Extraction and library preparation batches were randomized to include individuals from both species. Sequencing was performed on Illumina NovaSeqX, PE × 100 bp, with 39 samples sequenced in 2023 and the remaining 96 samples sequenced in 2025. Both species were represented in comparable numbers in each sequencing run (Suppl. Materials 1).

Average yield per sample was 3.05 Gb (± 0.76). Raw reads were processed according to an established bioinformatics pipeline (Carboni et al., 2025) (Table 1), obtaining an average of 7,410,484 (± 2,791,888) metagenomic pair-end reads per sample after removing low-quality reads, host contamination, and controls. We transferred the data into R software (R Core Team, 2024) and used the package decontam in prevalence mode with threshold 0.1 (Davis et al., 2018) to remove contaminants from the field and laboratory controls. For the taxonomic data, we retained only taxa present in more than 10 samples and with relative abundance higher than 0.01% (Parks et al., 2021), to improve precision and mitigate false-positive classifications associated with Kaiju (Edwin et al., 2024). It should be noted that no database currently exists specifically for the skin microbiota of non-human primates; therefore, species-level taxonomic assignments should always be interpreted with caution, as sequences may be assigned to closely related taxa. For functional analyses, we used MetaCyc pathway (PWY) abundance profiles and further classified them into broader functional categories using the microeco package (Liu et al., 2021). We estimated alpha diversity by calculating the Shannon index and richness using the phyloseq (McMurdie & Holmes, 2013) and vegan (Oksanen et al., 2024) packages. Beta diversity was assessed by calculating the Aitchison distance metric from CLR-transformed data, which accounts for the compositional nature of microbiome datasets (Gloor et al., 2017).

**Table 1.**
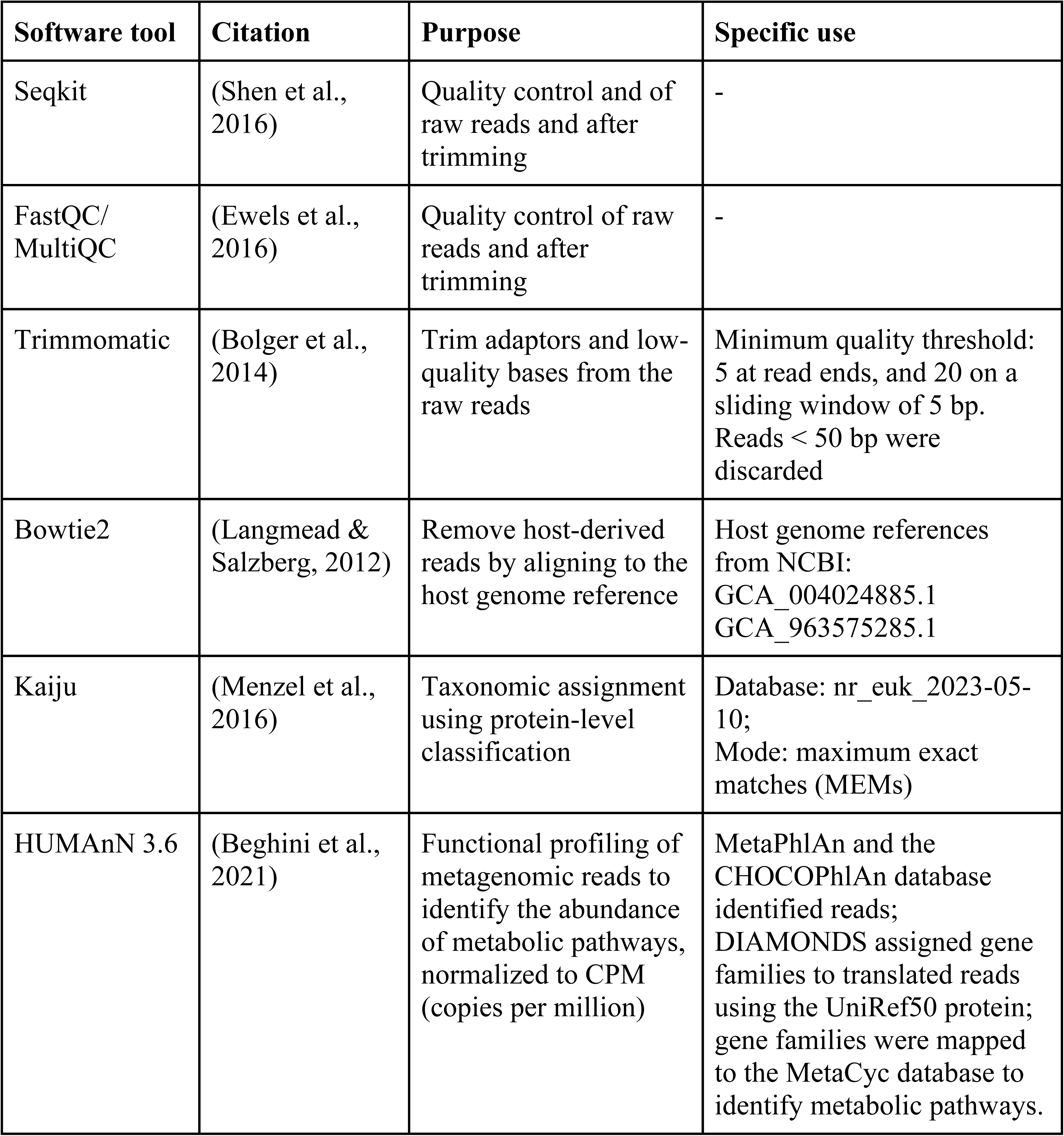
Bioinformatics tools used for microbiome analyses. The “Purpose” column reports the normal use of the tool, while in the column “Specific use” we report any specific use for our study.

| Software tool | Citation | Purpose | Specific use |
| --- | --- | --- | --- |
| Seqkit | (Shen et al., 2016) | Quality control and of raw reads and after trimming | - |
| FastQC/<br>MultiQC | (Ewels et al., 2016) | Quality control of raw reads and after trimming | - |
| Trimmomatic | (Bolger et al., 2014) | Trim adaptors and low-quality bases from the raw reads | Minimum quality threshold: 5 at read ends, and 20 on a sliding window of 5 bp. Reads < 50 bp were discarded |
| Bowtie2 | (Langmead & Salzberg, 2012) | Remove host-derived reads by aligning to the host genome reference | Host genome references from NCBI:<br>GCA_004024885.1<br>GCA_963575285.1 |
| Kaiju | (Menzel et al., 2016) | Taxonomic assignment using protein-level classification | Database: nr_euk_2023-05-10;<br>Mode: maximum exact matches (MEMs) |
| HUMAnN 3.6 | (Beghini et al., 2021) | Functional profiling of metagenomic reads to identify the abundance of metabolic pathways, normalized to CPM (copies per million) | MetaPhlAn and the CHOCOPhlAn database identified reads; DIAMONDS assigned gene families to translated reads using the UniRef50 protein; gene families were mapped to the MetaCyc database to identify metabolic pathways. |

### VOC analysis

We shipped the VOC samples annually on ice to the German Centre for Integrative Biodiversity Research (iDiv) in Leipzig, Germany, for chemical analyses via thermal desorption–gas chromatography-mass spectrometry (TD–GC-MS) using a Shimadzu QP2020 NX GC-MS linked to a TD-30R thermal desorption unit. Details on GC-MS parameters, as well as examples of chemical profiles, are provided in Suppl. Materials 2 - Table 1. We achieved chemical data interpretation using the software AMDIS 2.73 (Stein, 1999) and GC-MS Solutions v.4.20 (Shimadzu, Kyoto, Japan), and VOC identification using the NIST 14 mass spectral library (National Institute of Standards and Technologies, Gaithersburg, MD, USA), following methods detailed elsewhere (Poirier et al., 2025) (Suppl. Materials 2 - Table 1). We obtained a final list of 59 VOCs included in the analysis (Suppl. Materials 2 - Table 2). For each VOC in each odor profile, we computed relative peak abundance (RPA) = peak area/sum of peak areas in sample × 100 (excluding contaminants) and applied a log(x + 1)-transformation (hereafter logRPA) to centre the data distribution and to reduce the relative impact of abundant compounds while maintaining relative differences in the abundance of compounds within a sample. We further calculated total odor intensity as the log-transformed sum of all peak areas per profile. Finally, we estimated alpha diversity (Shannon index on RPA values) and beta diversity (Aitchison distances on CLR-transformed RPA values) for VOC profiles using the same tools as for the microbiome data.

**Table 2.**
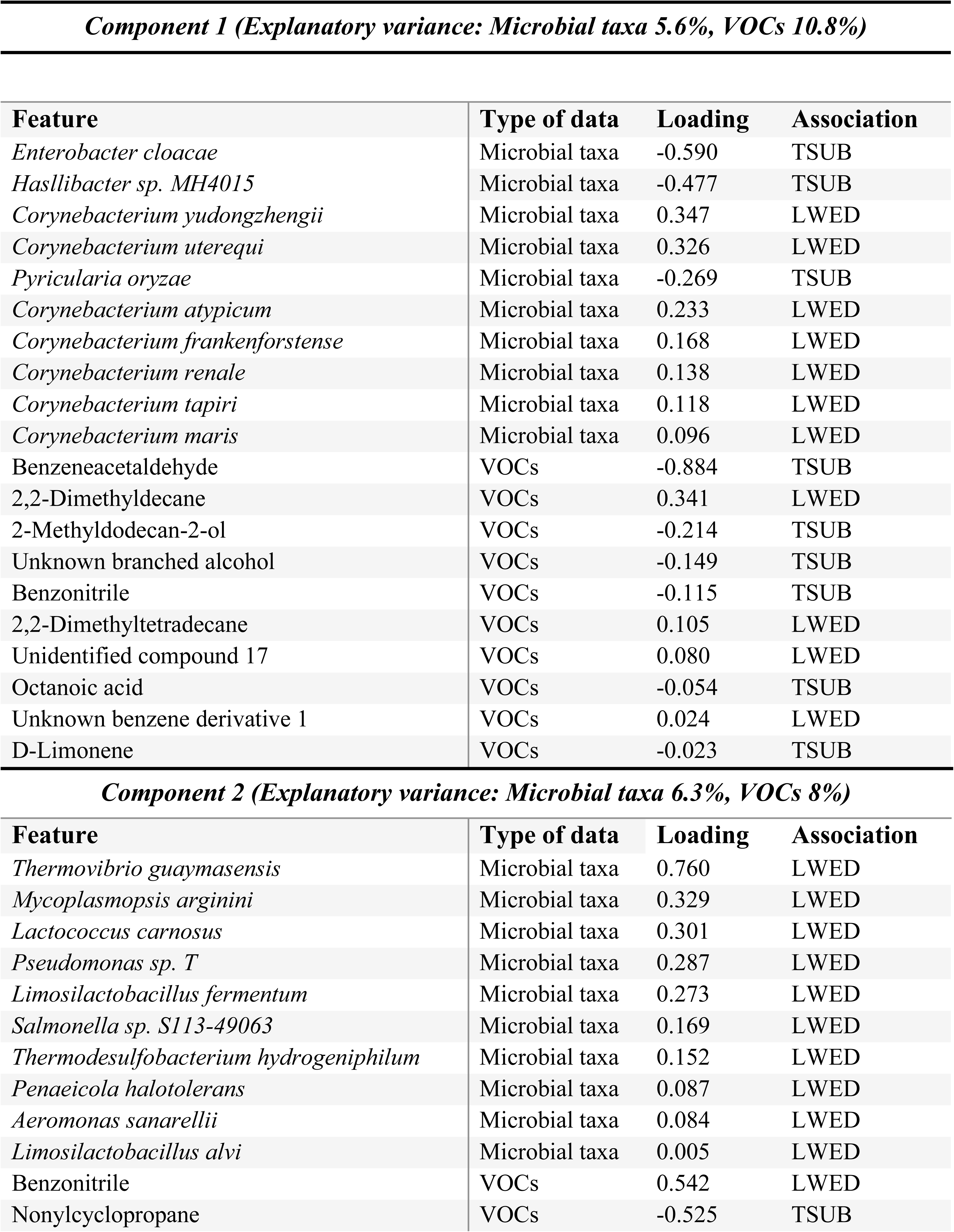
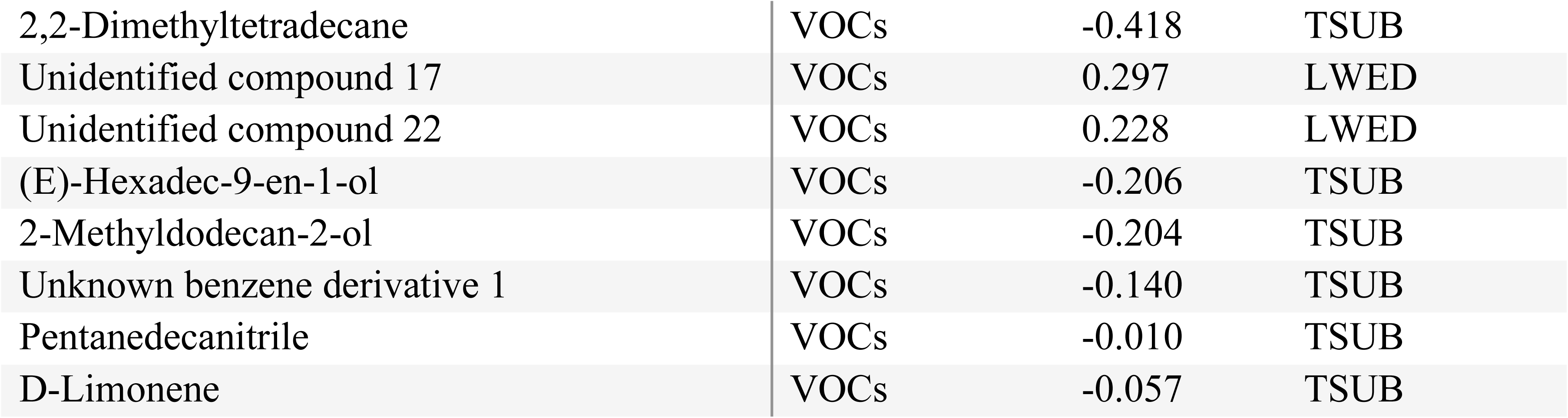
Top 20 features (i.e., microbial taxa and volatile organic compounds) with the highest absolute loading values, and therefore the strongest contributions to Components 1 and 2 of the multiblock-sPLS-DA integrating microbial taxonomic composition and odor profiles from the two tamarin species. Loading values indicate both the magnitude and direction of each feature’s contribution to species separation along the latent components. Positive loadings indicate features associated with *Leontocebus weddelli* (LWED) and negative loadings indicate features associated with *Tamarinus subgrisescens* (TSUB).

| **Component 1 (Explanatory variance: Microbial taxa 5.6%, VOCs 10.8%)** |  |  |  |
| --- | --- | --- | --- |
| **Feature** | **Type of data** | **Loading** | **Association** |
| *Enterobacter cloacae* | Microbial taxa | -0.590 | TSUB |
| *Hasllibacter sp. MH4015* | Microbial taxa | -0.477 | TSUB |
| *Corynebacterium yudongzhengii* | Microbial taxa | 0.347 | LWED |
| *Corynebacterium uterequi* | Microbial taxa | 0.326 | LWED |
| *Pyricularia oryzae* | Microbial taxa | -0.269 | TSUB |
| *Corynebacterium atypicum* | Microbial taxa | 0.233 | LWED |
| *Corynebacterium frankenforstense* | Microbial taxa | 0.168 | LWED |
| *Corynebacterium renale* | Microbial taxa | 0.138 | LWED |
| *Corynebacterium tapiri* | Microbial taxa | 0.118 | LWED |
| *Corynebacterium maris* | Microbial taxa | 0.096 | LWED |
| Benzeneacetaldehyde | VOCs | -0.884 | TSUB |
| 2,2-Dimethyldecane | VOCs | 0.341 | LWED |
| 2-Methyldodecan-2-ol | VOCs | -0.214 | TSUB |
| Unknown branched alcohol | VOCs | -0.149 | TSUB |
| Benzonitrile | VOCs | -0.115 | TSUB |
| 2,2-Dimethyltetradecane | VOCs | 0.105 | LWED |
| Unidentified compound 17 | VOCs | 0.080 | LWED |
| Octanoic acid | VOCs | -0.054 | TSUB |
| Unknown benzene derivative 1 | VOCs | 0.024 | LWED |
| D-Limonene | VOCs | -0.023 | TSUB |
| **Component 2 (Explanatory variance: Microbial taxa 6.3%, VOCs 8%)** |  |  |  |
| **Feature** | **Type of data** | **Loading** | **Association** |
| *Thermovibrio guaymasensis* | Microbial taxa | 0.760 | LWED |
| *Mycoplasmopsis arginini* | Microbial taxa | 0.329 | LWED |
| *Lactococcus carnosus* | Microbial taxa | 0.301 | LWED |
| *Pseudomonas sp. T* | Microbial taxa | 0.287 | LWED |
| *Limosilactobacillus fermentum* | Microbial taxa | 0.273 | LWED |
| *Salmonella sp. S113-49063* | Microbial taxa | 0.169 | LWED |
| *Thermodesulfobacterium hydrogeniphilum* | Microbial taxa | 0.152 | LWED |
| *Penaeicola halotolerans* | Microbial taxa | 0.087 | LWED |
| *Aeromonas sanarellii* | Microbial taxa | 0.084 | LWED |
| *Limosilactobacillus alvi* | Microbial taxa | 0.005 | LWED |
| Benzonitrile | VOCs | 0.542 | LWED |
| Nonylcyclopropane | VOCs | -0.525 | TSUB |
| 2,2-Dimethyltetradecane | VOCs | -0.418 | TSUB |
| Unidentified compound 17 | VOCs | 0.297 | LWED |
| Unidentified compound 22 | VOCs | 0.228 | LWED |
| (E)-Hexadec-9-en-1-ol | VOCs | -0.206 | TSUB |
| 2-Methyldodecan-2-ol | VOCs | -0.204 | TSUB |
| Unknown benzene derivative 1 | VOCs | -0.140 | TSUB |
| Pentanedecanitrile | VOCs | -0.010 | TSUB |
| D-Limonene | VOCs | -0.057 | TSUB |

### Statistical analyses

We conducted all statistical analyses in R (R Core Team, 2024). The final datasets included 135 samples for microbial taxonomy and metabolic pathways (*L. weddelli* n = 76, *T. subgrisescens* n = 59), and 134 odor profiles (*L. weddelli* n = 72, *T. subgrisescens* n = 62). The intersection of the microbiome and odor datasets included 114 samples (*L. weddelli* n = 60, *T. subgrisescens* n = 54). We first evaluated the degree of functional redundancy within the microbial community to assess the suitability of microbial metabolic pathway profiles for downstream analyses. Samples with higher microbial taxonomic diversity exhibited higher microbial pathway diversity (Spearman’s rank correlations: Shannon diversity, ρ = 0.300, p < 0.001; Richness, ρ = 0.621, p < 0.001) (Figure 1A, B) and taxonomic and functional community compositions were significantly associated (Mantel test: r = 0.246, p < 0.001, Procrustes test: r = 0.734, m² = 0.462, p = 0.001) (Figure 1C, D). The variation in functional composition suggested that functional redundancy was not strict in our dataset, and we therefore retained metabolic pathway profiles for subsequent analyses.

### Aim 1) the role of gland microbiomes and VOCs in mediating inter-specific discrimination

We tested our first aim using integrated and individual datasets. First, we integrated VOCs with microbial taxonomy and with microbial metabolic pathways, respectively, to evaluate their joint contribution to tamarin species differentiation. We performed two multiblock sPLS-DA analyses (i.e., DIABLO) using the *mixOmics* package (Rohart et al., 2017) on 114 samples, selecting two latent components per block. We first performed a PLS regression between each pair of CLR-normalized datasets, and used the resulting correlations to construct a design matrix with association weights of 0.55 between blocks. We then selected the number of variables to retain per component using *tune.block.splsda()*. For the taxonomy–VOC model, we retained 10 variables per block for both components; for the pathways–VOC model, we retained 15 and 10 pathway variables and 5 and 10 VOC variables across the two components. We cross-validated the models using the *perf()* M-fold function (5 folds, 50 repeats), and reported the AUC values as a measure of how well the integration of datasets discriminated between tamarin species. When repeating the same model separately for each sequencing run to assess potential batch effects, we obtained similar results, with consistent species separation.

**Figure 1.**
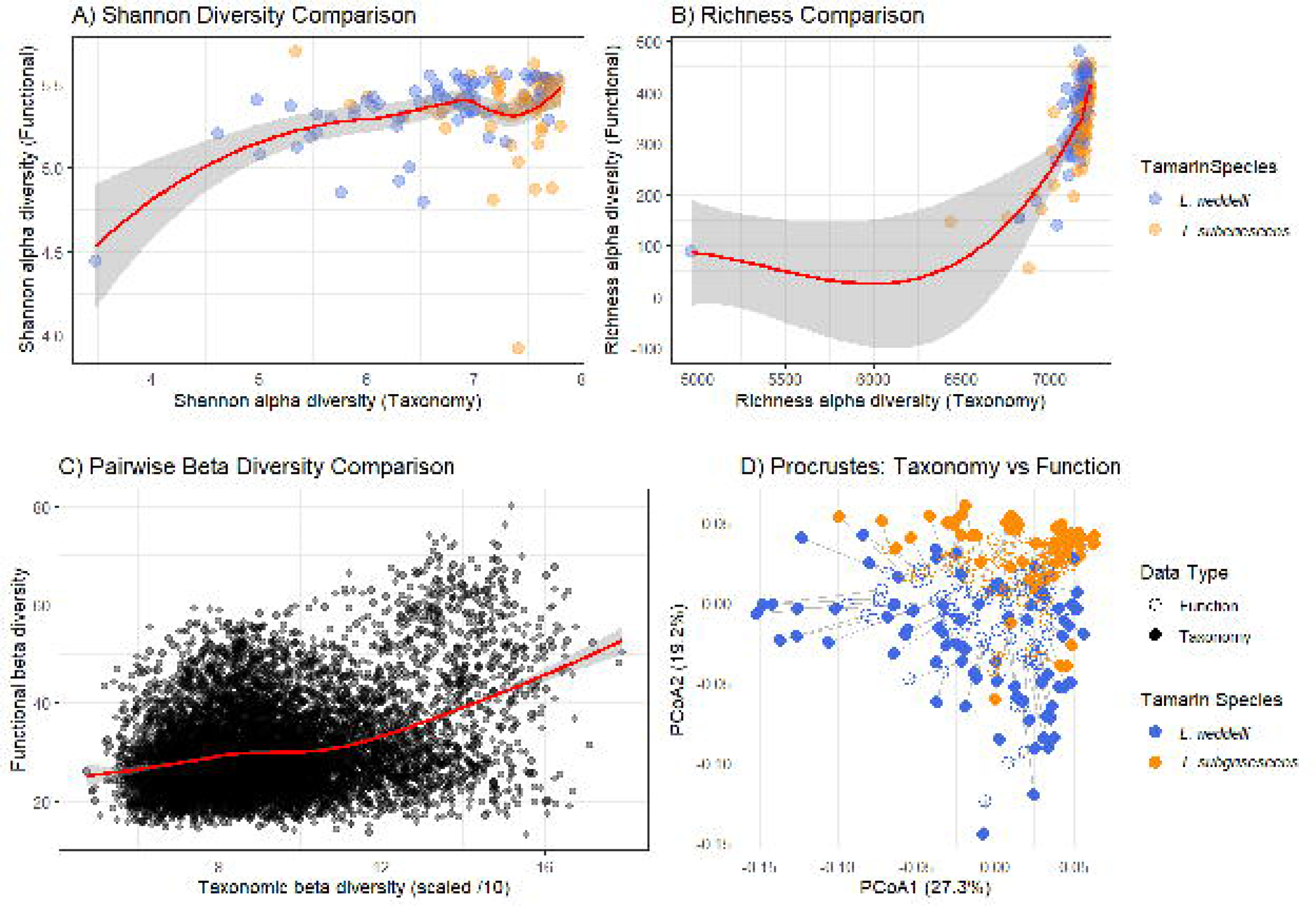
Panel **A)** compares the taxonomic (x-axis) and functional (y-axis) Shannon diversity across samples of both tamarin species, while Panel **B)** shows the same but for Richness index. Panel **C)** represents the comparison of pairwise taxonomic (x-axis) and functional (y-axis) Aitchison diversity among samples. Taxonomic beta diversity values were scaled by a factor of 10 for visualization. In these three panels, red lines indicate loess-smoothed trends with 95% confidence intervals. Panel **D)** shows the Procrustes analysis comparing taxonomic and functional community ordinations. Segments connect the positions of the same sample in taxonomic and functional ordination spaces; shorter arrows indicate greater concordance between taxonomic composition and functional potential.

We then analyzed the three datasets separately by testing species differences in alpha diversity, beta diversity, and feature abundances. We fitted six linear models to analyze alpha diversity metrics: generalized linear mixed models for the microbiome datasets (*glmer()*, lme4 package) and linear mixed models for the VOC dataset (*lmer()*, lme4 package) (Bates et al., 2015) (Suppl. Materials 3 - Tables 1-3). Sequencing run was included to account for differences between sequencing batches, and read count was included to control for variation in sequencing depth because samples were not rarefied to avoid data loss. We also included year of collection to account for batch variation among samples, and capture day (13–22 capture days per season) to control for potential methodological shifts during sample collection. We assessed model performance by comparing AICc values between the full models and their corresponding null models. We assessed model fit and assumptions using the DHARMa package (Hartig et al., 2024). We found no evidence of multicollinearity (*vif()* in the car package) (Fox & Weisberg, 2019), and residuals showed no evidence of autocorrelation, with Durbin–Watson statistics close to 2. We then tested for differences in Aitchison distances in the two microbiome datasets by performing PERMANOVA analyses using the vegan package (Oksanen et al., 2024), assessing the effect of tamarin species while controlling for sequencing run and stratifying by year of collection, thereby accounting for individuals sampled repeatedly across different years. For the VOC dataset, variance between species was heterogeneous, violating PERMANOVA assumptions. We therefore used a redundancy analysis (RDA) controlling for capture day and stratifying by year of collection (Suppl. Materials 3 - Tables 4,5). Finally, we used the ancombc2 package (Lin & Peddada, 2024) (“holm” method) to identify differentially abundant microbial taxa and metabolic pathways between tamarin species, and Wilcoxon rank-sum tests with Holm-adjusted p-values to assess species differences in the abundance of individual VOCs.

### Aim 2) the role of gland microbiomes and VOCs in communicating group membership

We tested our second aim by considering the two tamarin species separately. We ran similar tests as above. For alpha diversity, we fitted twelve linear models (Suppl. Materials 3 - Tables 6-11) and evaluated model performance and assumptions using the same procedures described above. Capture day was not included in the VOC models because it was collinear with group identity, as entire tamarin groups were typically captured during a single capture event. Because multiple groups were compared, pairwise differences were assessed using estimated marginal means with Tukey-adjusted post hoc tests (Lenth & Piaskowski, 2026). For beta diversity, we applied the same framework as above, using group membership as the main predictor in both PERMANOVA and RDA models (Suppl. Materials 3 - Tables 12,13). We also assessed within-species differences in the abundance of microbial taxa and metabolic pathways among groups using ANCOM-BC2 (“Holm” method), and differences in VOC abundance using Wilcoxon rank-sum tests (“Holm” correction).

## RESULTS

### i) Multiomics species-specific discrimination

Scent gland microbiota of both tamarin species were dominated by bacterial taxa (*L. weddelli:* 78.0%, *T. subgrisescens:* 65.2% of total reads), followed by eukaryotes (*L. weddelli*: 21.9%, *T. subgrisescens:* 34.7%) (Figure 2A). The genera with the highest mean relative abundance in *L. weddelli* were *Bifidobacterium* (13.2%), *Staphylococcus* (11.0%), and *Corynebacterium* (3.6%), whereas in *T. subgrisescens* the most abundant genera were *Malassezia* (4.8%), *Bifidobacterium* (4.3%), and *Actinomycetospora* (3.7%). In both species, the most abundant metabolic pathways were related to amino acid and nucleotide biosynthesis, fermentation, carbohydrate, lipid, and fatty acid biosynthesis (Figure 2B, C). For odor profiles, we retrieved 59 VOCs (mean ± SD = 5 ± 2 per sample), and we identified 61% of these, including hydrocarbons (e.g., aliphatic and cyclic alkanes, 36.4 %), fatty acids (18.2 %), nitrogen-containing compounds (e.g., amides, nitriles, amines, 18.2 %), alcohols (12.1 %), and aldehydes (9.1 %) (Suppl. Materials 2 - Table 2). The most prevalent identified VOCs in *L. weddelli* were benzeneacetaldehyde (in 36.1 % of samples), nonylcyclopropane (34.7 %), pentadecanenitrile (29.2 %) and 2,2-dimethyltetradecane (27.8 %), and in *T. subgrisescens* were pentadecanenitrile (33.9 %), 2,2-dimethyltetradecane (29.0 %) and nonylcyclopropane (25.8 %) (Suppl. Materials 2 - Table 2).

**Figure 2.**
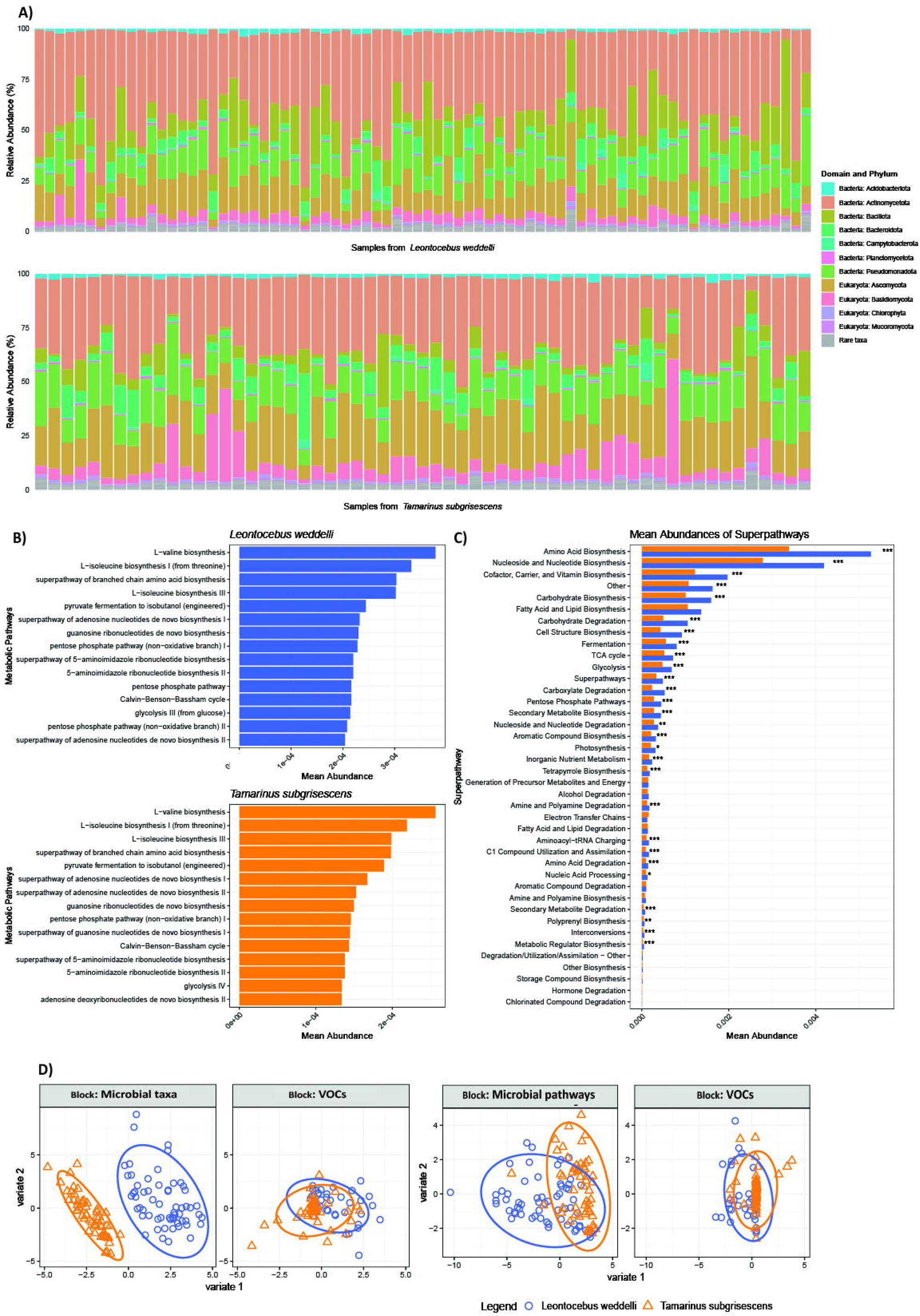
Panel **A)** shows the relative abundance of microbial taxa at the phylum level from suprapubic glands of *L. weddelli* (n = 76) and *T. subgrisescens* (n = 59). The legend only shows the 11 phyla with mean abundance > 0.005, while the rest are considered “Rare taxa”, in grey. Panel **B)** displays the relative abundance (copies per million - CPM) of the 15 most abundant metabolic pathways from the same samples. Panel **C)** shows the relative abundance (CPM) of microbial metabolic super-pathways in the two tamarin species. Super-pathways group together all the metabolic pathways with the same border functional role. Pathways that could not be assigned to a super-pathway category were grouped under “Other”. Asterisks denote significant differences in abundance between tamarin species (Wilcoxon rank-sum tests, Holm-adjusted p-values). Panel **D)** displays the results of multiblock-sPLS-DA analyses performed on samples from *L. weddelli* (n = 60) and *T. subgrisescens* (n = 54). The two blocks on the left show results from the model integrating microbial taxonomy with odor profiles, while the two blocks on the right show results from the model integrating microbial metabolic pathways with odor profiles. Each point represents an individual sample positioned on the first two latent components. Ellipses represent 90% confidence intervals and greater separation indicates that the integrated datasets discriminate strongly between tamarin species.

The integration of microbial taxa and VOC profiles contributed to the discrimination between tamarin species, with taxonomic composition showing stronger discriminatory power than VOC composition (Figure 2D). Cross-validation analyses supported this pattern (Component 1: AUC = 0.85, p = 0.001; Component 2: AUC = 0.87, p < 0.001), with the microbial taxa block showing higher accuracy compared to the VOC block (Suppl. Materials 3 - Table 14). The microbial taxa contributing more strongly to species discrimination were bacteria from the genera *Enterobacter* and *Corynebacterium*, while the VOCs included mostly aromatic compounds and alkanes, especially benzeneacetaldehyde, benzonitrile, 2,2-dimethyldecane, and nonylcyclopropane (Table 2). Benzonitrile and nonylcyclopropane were also among the VOCs correlating the most with the microbial taxa with the highest loadings (Figure 3A).

The integration of microbial metabolic pathways and VOC matrices also contributed to species discrimination (Figure 2D) (Component 1: AUC = 0.79, p = 0.004; Component 2: AUC = 0.83, p < 0.001), with higher accuracy for the metabolic pathways (Suppl. Materials 3 - Table 14). The pathways related to cell wall biosynthesis, formaldehyde assimilation, pyrimidine and chitin metabolism strongly contributed to species discrimination, while, as expected, results for the VOCs were similar to the previous model (Table 3). Interestingly, selected VOCs showed stronger correlations with the metabolic pathways than with taxa overall (Figure 3B).

**Figure 3.**
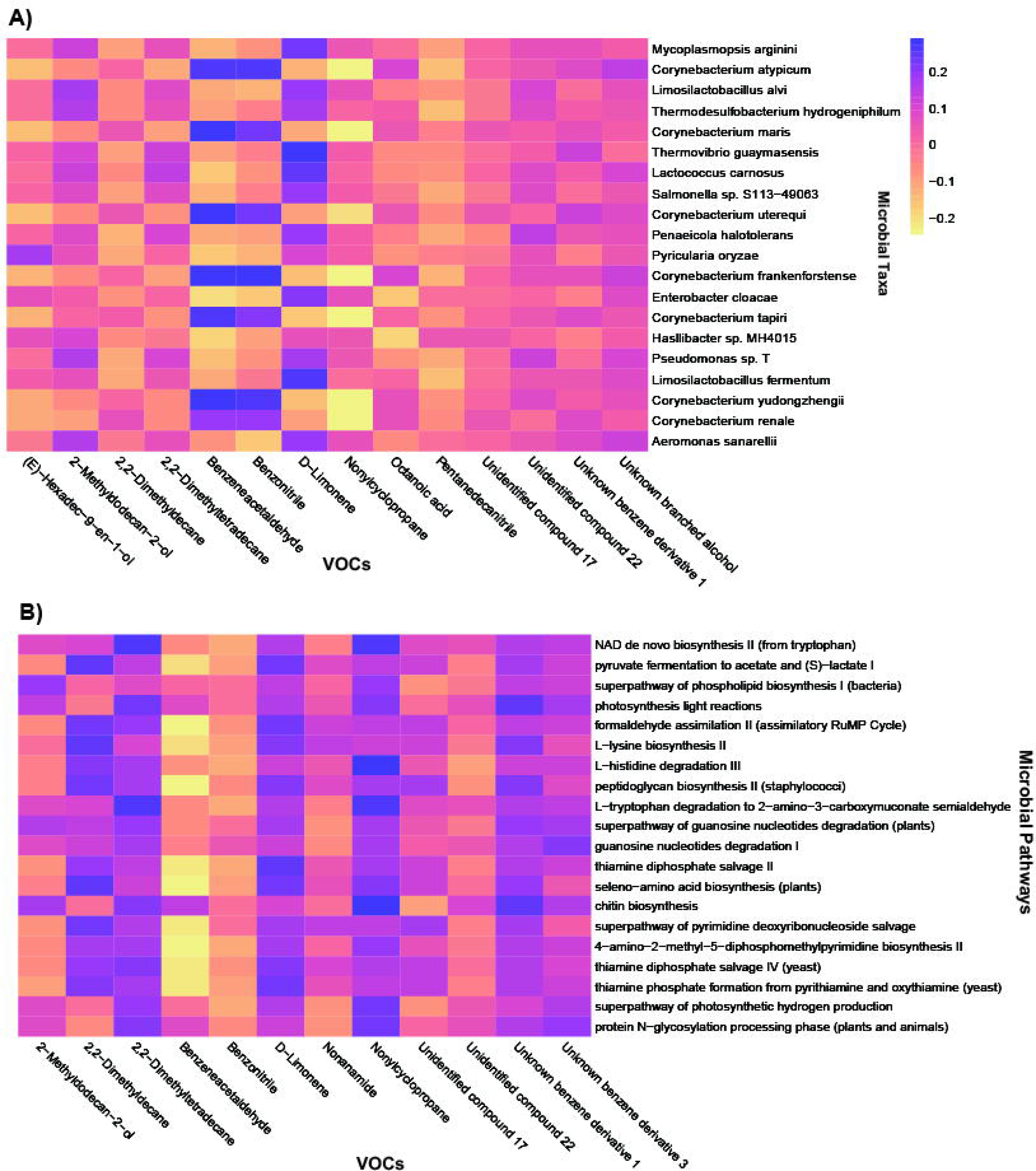
Panel **A)** represents a Spearman correlation heatmap between the microbial taxa and VOCs with the highest absolute loading values across Components 1 and 2 of the multiblock-sPLS-DA model integrating microbial taxonomy and odor components. Panel **B)** shows a Spearman correlation heatmap between the microbial metabolic pathways and VOCs with the highest absolute loading values across Components 1 and 2 of the multiblock-sPLS-DA model integrating microbial metabolic pathways and odor components. The selected features are those contributing most strongly to discrimination between the two tamarin species. Heatmap colors indicate the strength and direction of the correlation: darker colors representing positive correlations and lighter colors representing negative correlations.

**Table 3.**
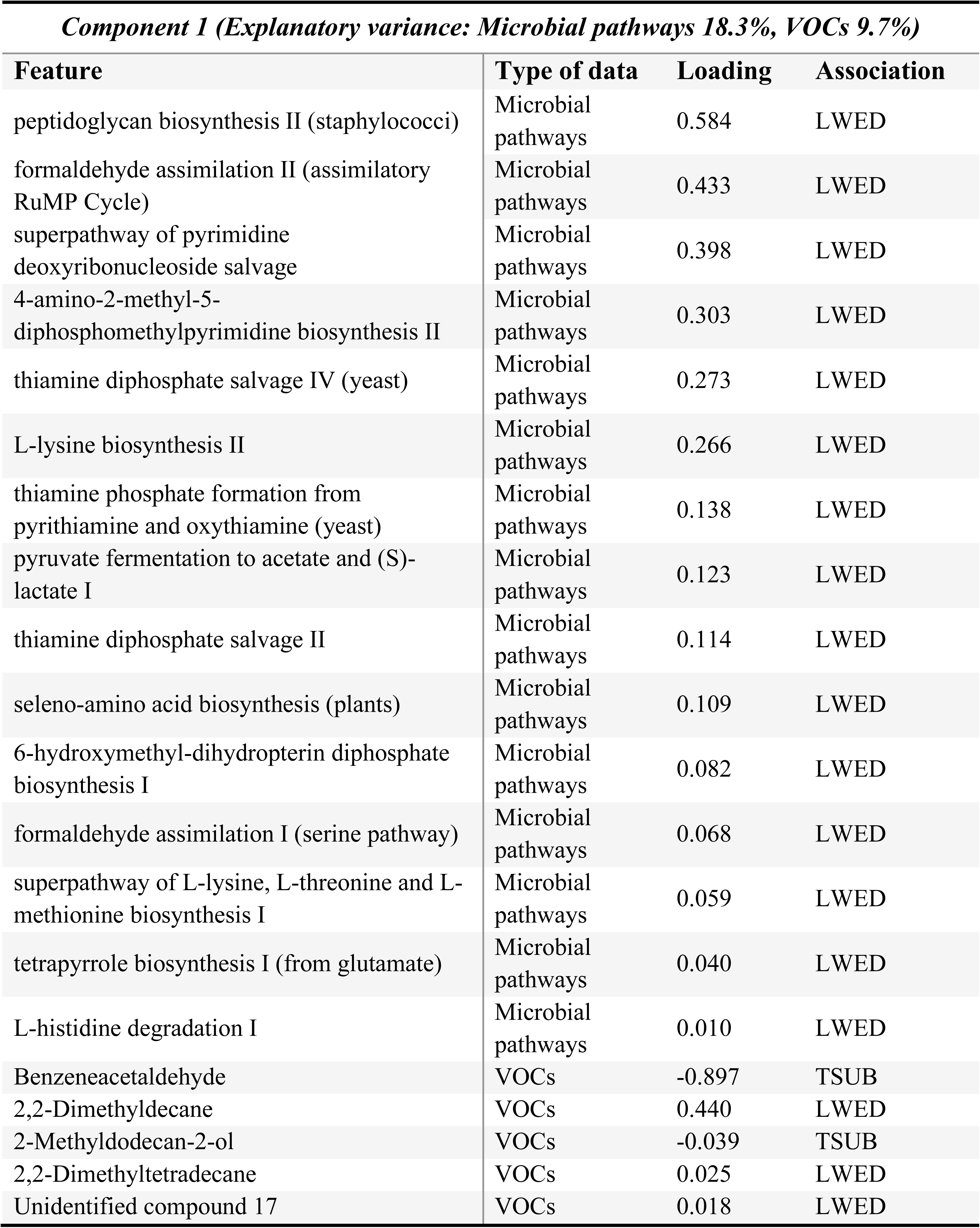
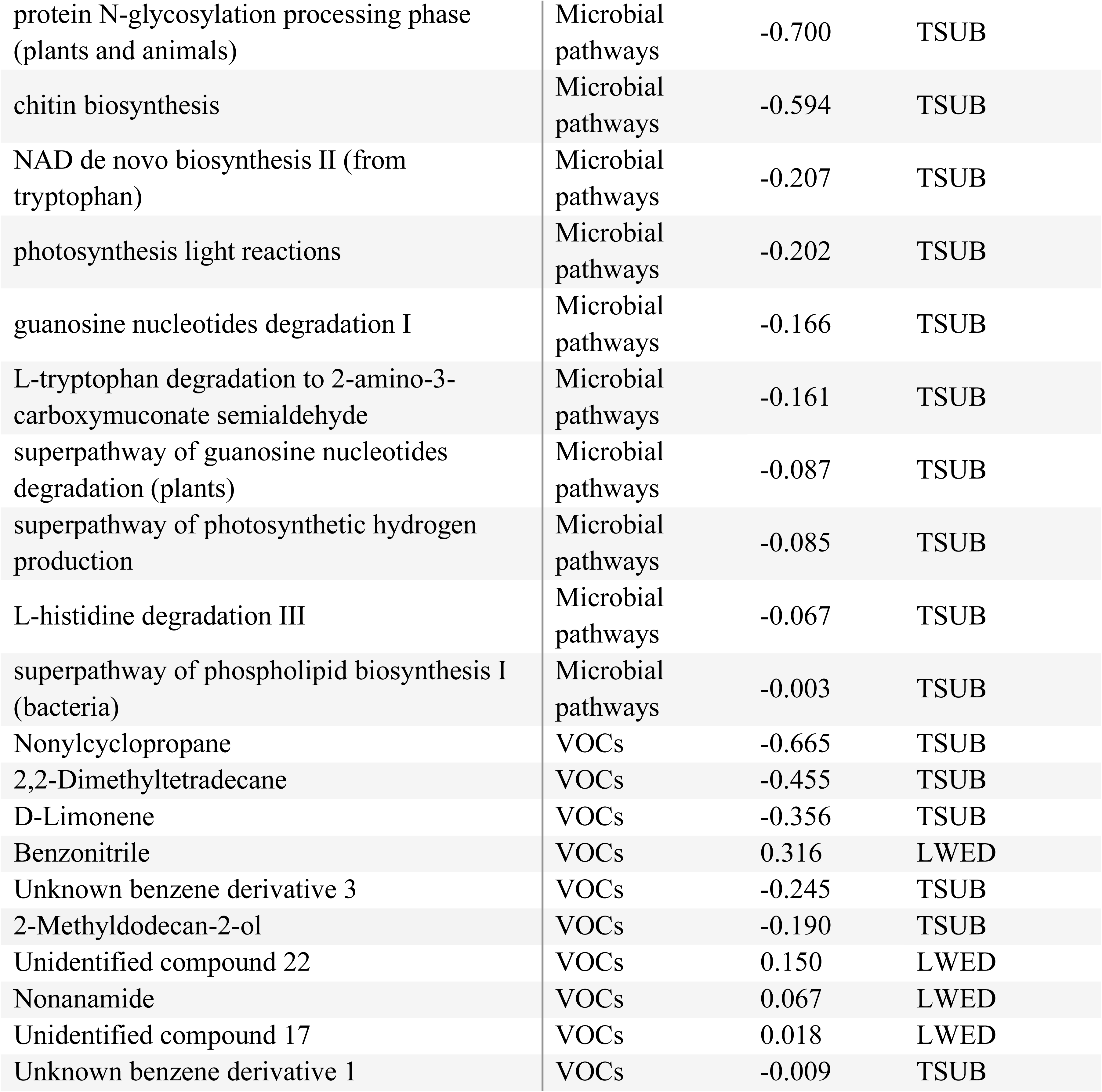
Top 20 features (i.e., microbial metabolic pathways and volatile organic compounds) with the highest absolute loading values, and therefore the strongest contributions to Components 1 and 2 of the multiblock-sPLS-DA integrating microbial metabolic profile and odor profile from the two tamarin species. Loading values indicate both the magnitude and direction of each feature’s contribution to species separation along the latent components. Positive loadings indicate features associated with *Leontocebus weddelli* (LWED) and negative loadings indicate features associated with *Tamarinus subgrisescens* (TSUB).

| *Component 1 (Explanatory variance: Microbial pathways 18.3%, VOCs 9.7%)* |  |  |  |
| --- | --- | --- | --- |
| Feature | Type of data | Loading | Association |
| peptidoglycan biosynthesis II (staphylococci) | Microbial pathways | 0.584 | LWED |
| formaldehyde assimilation II (assimilatory RuMP Cycle) | Microbial pathways | 0.433 | LWED |
| superpathway of pyrimidine deoxyribonucleoside salvage | Microbial pathways | 0.398 | LWED |
| 4-amino-2-methyl-5-diphosphomethylpyrimidine biosynthesis II | Microbial pathways | 0.303 | LWED |
| thiamine diphosphate salvage IV (yeast) | Microbial pathways | 0.273 | LWED |
| L-lysine biosynthesis II | Microbial pathways | 0.266 | LWED |
| thiamine phosphate formation from pyrithiamine and oxythiamine (yeast) | Microbial pathways | 0.138 | LWED |
| pyruvate fermentation to acetate and (S)-lactate I | Microbial pathways | 0.123 | LWED |
| thiamine diphosphate salvage II | Microbial pathways | 0.114 | LWED |
| seleno-amino acid biosynthesis (plants) | Microbial pathways | 0.109 | LWED |
| 6-hydroxymethyl-dihydropterin diphosphate biosynthesis I | Microbial pathways | 0.082 | LWED |
| formaldehyde assimilation I (serine pathway) | Microbial pathways | 0.068 | LWED |
| superpathway of L-lysine, L-threonine and L-methionine biosynthesis I | Microbial pathways | 0.059 | LWED |
| tetrapyrrole biosynthesis I (from glutamate) | Microbial pathways | 0.040 | LWED |
| L-histidine degradation I | Microbial pathways | 0.010 | LWED |
| Benzeneacetaldehyde | VOCs | -0.897 | TSUB |
| 2,2-Dimethyldecane | VOCs | 0.440 | LWED |
| 2-Methyldodecan-2-ol | VOCs | -0.039 | TSUB |
| 2,2-Dimethyltetradecane | VOCs | 0.025 | LWED |
| Unidentified compound 17 | VOCs | 0.018 | LWED |

**Component 2 (Explanatory variance: Microbial pathways 6.2%, VOCs 8.1%)**
| **Feature** | **Type of data** | **Loading** | **Association** |
| --- | --- | --- | --- |
| protein N-glycosylation processing phase (plants and animals) | Microbial pathways | -0.700 | TSUB |
| chitin biosynthesis | Microbial pathways | -0.594 | TSUB |
| NAD de novo biosynthesis II (from tryptophan) | Microbial pathways | -0.207 | TSUB |
| photosynthesis light reactions | Microbial pathways | -0.202 | TSUB |
| guanosine nucleotides degradation I | Microbial pathways | -0.166 | TSUB |
| L-tryptophan degradation to 2-amino-3-carboxymuconate semialdehyde | Microbial pathways | -0.161 | TSUB |
| superpathway of guanosine nucleotides degradation (plants) | Microbial pathways | -0.087 | TSUB |
| superpathway of photosynthetic hydrogen production | Microbial pathways | -0.085 | TSUB |
| L-histidine degradation III | Microbial pathways | -0.067 | TSUB |
| superpathway of phospholipid biosynthesis I (bacteria) | Microbial pathways | -0.003 | TSUB |
| Nonylcyclopropane | VOCs | -0.665 | TSUB |
| 2,2-Dimethyltetradecane | VOCs | -0.455 | TSUB |
| D-Limonene | VOCs | -0.356 | TSUB |
| Benzonitrile | VOCs | 0.316 | LWED |
| Unknown benzene derivative 3 | VOCs | -0.245 | TSUB |
| 2-Methyldodecan-2-ol | VOCs | -0.190 | TSUB |
| Unidentified compound 22 | VOCs | 0.150 | LWED |
| Nonanamide | VOCs | 0.067 | LWED |
| Unidentified compound 17 | VOCs | 0.018 | LWED |
| Unknown benzene derivative 1 | VOCs | -0.009 | TSUB |

When we analyzed the three datasets separately, tamarin species differed significantly in microbial taxonomic diversity and composition. *T. subgrisescens* had higher Shannon taxonomic diversity than *L. weddelli* (GLMM: Estimate = 0.083; p < 0.001), but comparable richness (Suppl. Materials 3 - Table 1). Tamarin species significantly explained 5.9% of taxonomic variation (PERMANOVA: F = 8.38, p = 0.001; Suppl. Materials 3 - Table 4) (Figure 4A), a pattern driven by *Corynebacterium frankfortense, Staphylococcus auricularis, Malassezia japonica* and *M. obtusa* (Figure 4B). However, diversity measures of metabolic pathways or VOCs did not differ between species (Suppl. Materials 3 - Tables 2-5), and null models outperformed alternative models for the Shannon diversity of both datasets. Differential abundance analyses for each dataset identified 198 microbial species enriched in *L. weddelli*, 130 in *T. subgrisescens* (Figure 4C), two pathways enriched in *L. weddelli* (L-phenylalanine degradation IV: LFC = -0.664; L-ascorbate biosynthesis IV: LFC = -0.567), while among the VOCs, 3,5-dimethylundecane, acetic acid and unknown amide and branched ester were exclusively detected in samples from *L. weddelli* (Suppl. Materials 2 - Figure 1).

**Figure 4.**
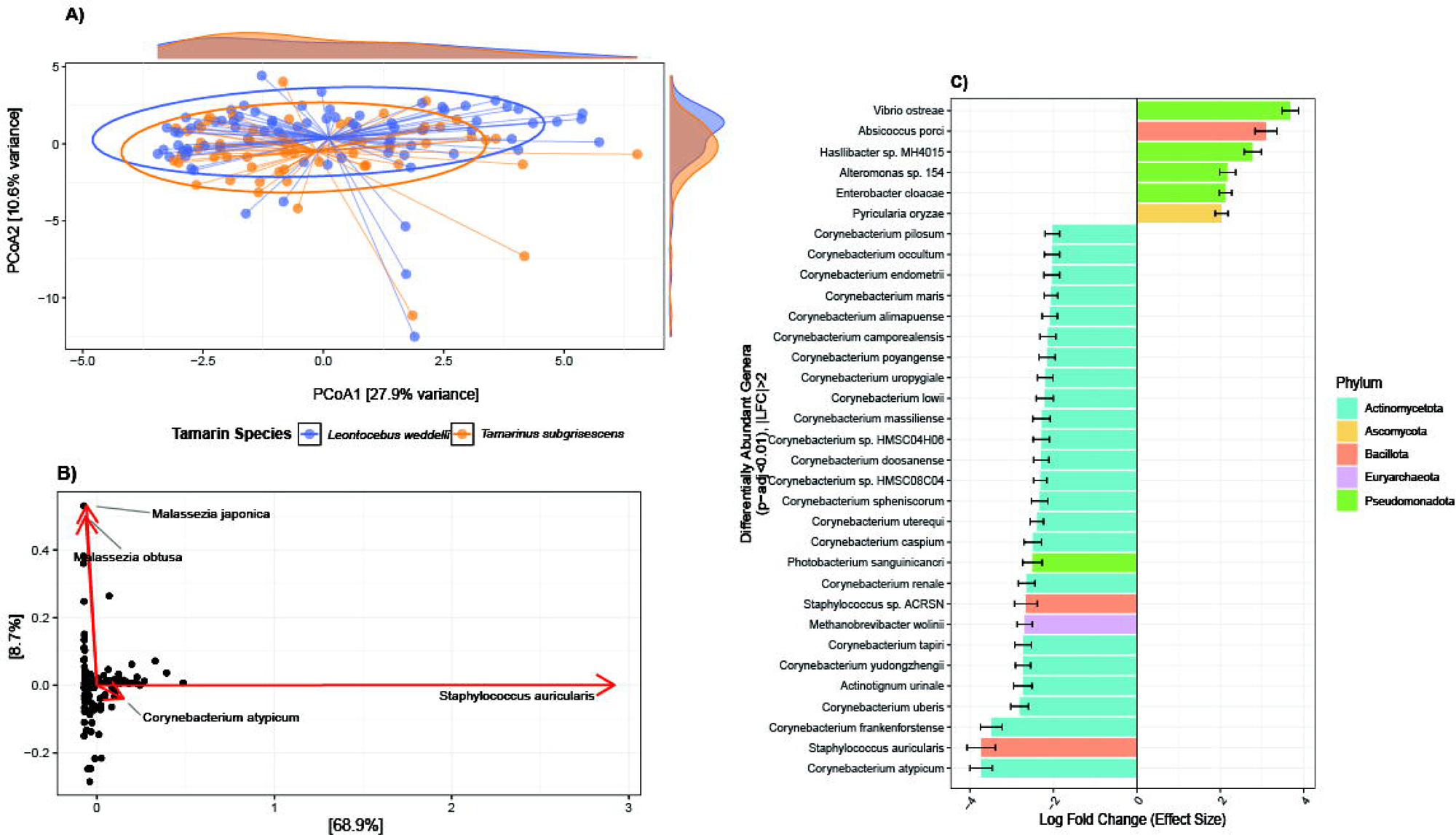
Panel **A)** shows the Principal Coordinates Analysis (PCoA) based on Aitchison distances showing variation in microbial community composition based on tamarin species. Ellipses represent the 85% confidence interval for each species, and marginal density plots show the distribution of samples along each axis. Panel **B)** shows the Principal Component Analysis (PCA) of CLR-transformed microbial taxonomic abundances. Red arrows indicate the taxa with the largest loadings on PC1 and PC2, which contribute the most to differences in community composition. Panel **C)** displays the results of the ANCOM-BC2 test on microbial taxa by tamarin species. Only taxa that passed sensitivity analysis with an absolute log fold change > 2, adjusted p < 0.01 are shown. Negative log fold changes indicate taxa enriched in *L. weddelli*, whereas positive values indicate taxa enriched in *T. subgrisescens*. The color of the bars represents the phylum of each microbial species.

### ii) Multiomics group-specific discrimination

Social groups within species did not differ in alpha diversity for microbial taxa, metabolic pathways, or VOCs for either tamarin species, and the models for the Shannon index of both microbial datasets and the VOC dataset failed to outperform their respective null models (Suppl. Materials 3 - Tables 6-11). However, group membership explained 12.4% of variation in beta diversity of the microbial taxonomy of *L. weddelli* (PERMANOVA: F = 1.67, p = 0.009, Suppl. Materials 3 - Table 12) (Figure 5A), and was a significant predictor of sample VOC composition in *T. subgrisescens* (RDA: F = 1.17, p = 0.036; Suppl. Materials 3 - Table 13) (Figure 5B). However, it was not a significant predictor for diversity of taxonomy of *T. subgrisescens*, metabolic pathways of both species, and VOCs for *L. weddelli* (Suppl. Materials 3 - Tables 12, 13). The abundance of specific microbial taxa varied across group dyads of both tamarin species (Figure 5C, D) and only in *T. subgrisescens* for the metabolic pathways (Figure 5E).

**Figure 5.**
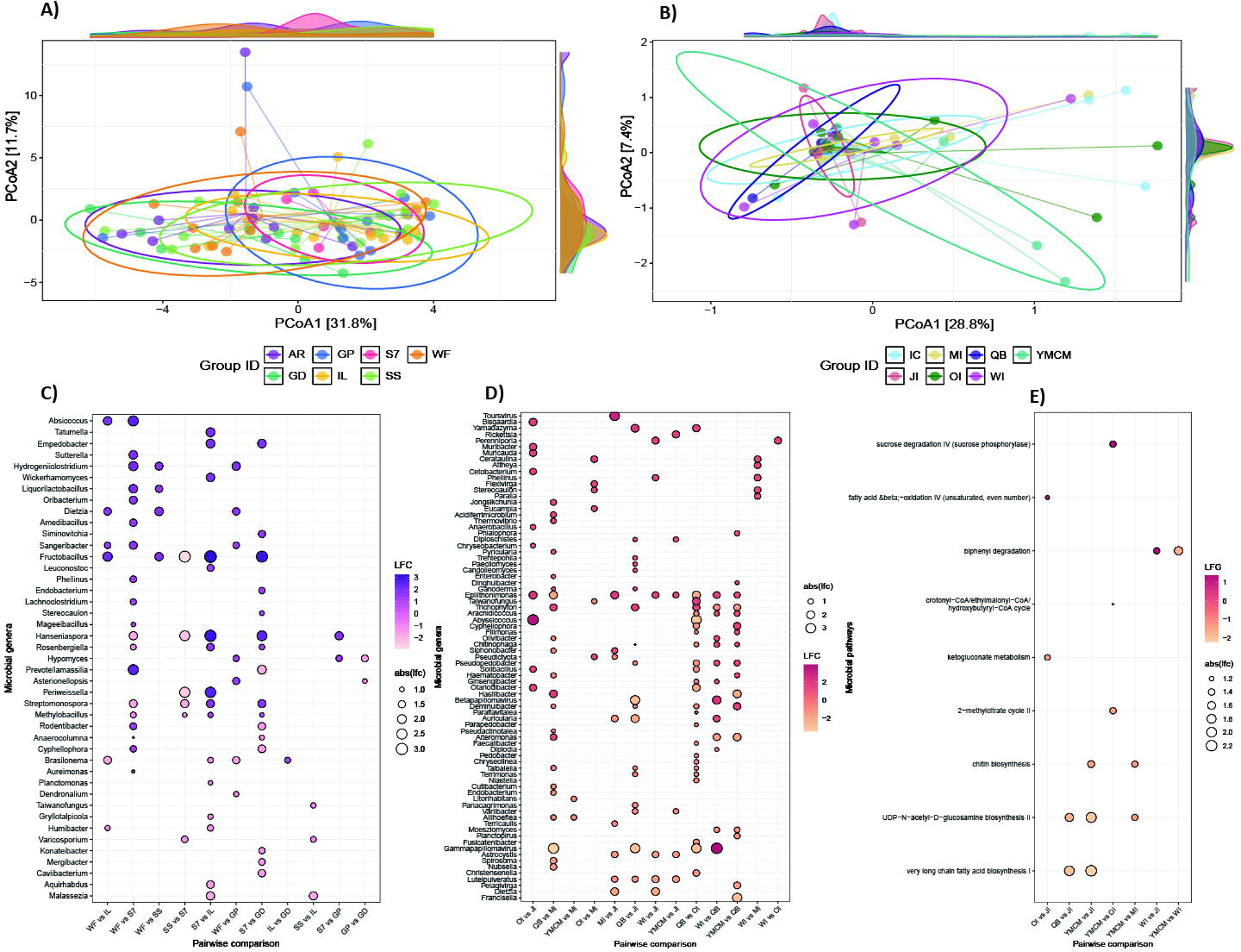
Panel **A)** and **B)** illustrate the PCoA based on Aitchison distances showing variation in microbial community composition based on group membership of *L. weddelli* (A), and in VOC composition based on group membership of *T. subgrisescens* (B). Ellipses represent the 85% confidence interval, and marginal density plots show the distribution of samples along each axis. Panel **C)** shows the results of the ANCOM-BC2 analysis comparing microbial taxonomic genera between pairs of social groups in *L. weddelli*. Positive log fold change (LFC) values indicate taxa enriched in the first group of each comparison, whereas negative values indicate taxa enriched in the second group. Panel **D)** shows the corresponding ANCOM-BC2 results for pairwise comparisons of microbial genera among social groups of *T. subgrisescens*. Panel **E)** presents the results of the ANCOM-BC2 analysis of microbial metabolic pathways among social groups of *T. subgrisescens*.

## DISCUSSION

Our study provides among the first integrative molecular investigations of the scent glands of wild sympatric monkeys, combining shotgun metagenomic and VOC metabolomic data to investigate the potential role of olfaction in maintaining reproductive isolation and social communication. Closely related sympatric tamarins differed in microbial taxonomy and their relationships with VOCs, in accordance with the hypothesis that olfaction may be contributing to species recognition and maintenance of reproductive isolation. In contrast, evidence for group-specific molecular signatures was limited and differed between species. Together, these findings indicate that scent gland microbiomes differ more between species than between social groups, although questions remain on how this reflects selection on olfactory communication.

### i) Multiomics species-specific discrimination

In accordance with our hypotheses, we found that the two wild sympatric species could be discriminated based on the covariation of odor and microbiome of their scent glands, although primarily based on microbial taxonomic composition. Despite their high potential for hybridization given similarities in behavior, physiology, and ecology (Coombes et al., 2018; M. Rowe et al., 2020; Zinner et al., 2011), these two tamarin species remain reproductively isolated, and the species-specific differences found here point to olfaction as a potential mechanism involved in maintaining this isolation. *Corynebacterium* spp. defined *L. weddelli*, while members of the *Proteobacteria* associated with *T. subgrisescens*. Although some aspects of microbial variation may reflect species-specific ecological filtering or skin physiology rather than selection for odor production alone, these taxa have been previously implicated in the production of volatile fatty acids, aldehydes, alcohols, and methyl esters involved in scent communication (Hou et al., 2024; Leclaire et al., 2017; Natsch et al., 2003). Their putative functional relevance was consistent with the association between VOC variation and microbial metabolic pathways involved in amino acid metabolism and fermentation. Furthermore, the VOCs that covaried with both microbial datasets included aromatic compounds such as benzeneacetaldehyde, benzonitrile, and D-limonene in *T. subgrisescens*, and methyl alkanes such as 2,2-dimethyldecane and 2,2-dimethyltetradecane in *L. weddelli*, many of which have been previously reported in mammalian scent (Kücklich et al., 2019; Medrano et al., 2024; Peng et al., 2010; Poirier et al., 2021a; Setchell et al., 2010).

Notably, in both integrative models, individuals of *T. subgrisescens* showed relatively little variation in VOC profiles along the first component (Figure 2D, variate 1), with a similar, although less pronounced, pattern observed for microbial taxonomy and predicted metabolic pathways. This suggests that odor and microbiome profiles may be more conserved in *T. subgrisescens* than in *L. weddelli*. Although these associations do not demonstrate causality, they are consistent with the hypothesis that more homogeneous microbial communities may contribute to more conserved odor profiles, highlighting the value of integrating microbial taxonomic, functional, and chemical data to generate hypotheses about mechanisms underlying odor variation and species discrimination.

When analyzed independently, microbial taxonomy revealed significant differences in diversity and community composition between the two tamarin species. Microbes are selected by both environmental exposure and host-mediated ecological filtering through immune defenses, gland physiology, and morphology (Foster et al., 2017; Zeng et al., 2017). Because our species occurred sympatrically and likely experienced similar environmental microbial pools, the observed differences may instead reflect variation in host characteristics that shape microbial establishment and persistence. Notably, *L. weddelli* showed lower Shannon diversity (but comparable richness) and increased relative abundance of *Corynebacterium* spp. compared to *T. subgrisescens*. Identifying the host-specific selective pressures that shaped the assembly of this distinctive microbial community may provide insights into sympatric species discrimination through gland microbiome and its functional consequences. For example, *Corynebacterium* may contribute to the scent of *L. weddelli* through its ability to metabolize host-derived lipids and amino acid derivatives into volatile fatty acid compounds (Lam et al., 2018; Natsch et al., 2003; Nordstrom et al., 1989; Rojas et al., 2023; Zhou et al., 2021). Interestingly, in this study acetic acid, a volatile fatty acid commonly detected in the scent secretions of primates (Gallagher et al., 2008; Morelli et al., 2014; Smith, 2001; Vaglio et al., 2016) and other mammals (Apps et al., 2012; Gassett et al., 1996; Hagey & MacDonald, 2003; Salamon & Davies, 1998; Weiß et al., 2018), was exclusively found in *L. weddelli*. It therefore warrants targeted investigation as a candidate compound contributing to the distinct scent profile of *L. weddelli* and the putative role of *Corynebacterium* in its production.

Predicted microbial metabolic pathways also supported the role of microbes in odor production, although species-specific differentiation was limited. This pattern is consistent with the functional redundancy hypothesis, as taxonomically distinct microbial communities retain similar metabolism as they perform essential ecological functions (Fässler et al., 2025). In our dataset, we found a strong Procrustes correlation, indicating similar overall multivariate structure between taxonomic and functional profiles, whereas pairwise distances were only weakly positively correlated in the Mantel test. This suggests that differences in taxonomic composition do not consistently translate into proportional differences in metabolic pathways among samples. Although the lower microbial taxonomic diversity observed in *L. weddelli* may reflect a smaller repertoire of genes, and thus greater functional specialization (Galand et al., 2018), this pattern could have been obscured by the substantial overlap in core metabolic functions. Future experimental analyses of key microbial taxa and their metabolic activity could help identify the pathways directly involved in odor production.

Although our VOC analysis did not reveal significant species-specific differences in overall VOC diversity or composition, a few compounds, namely 3,5-dimethylundecane, acetic acid, an unknown amide, and an unknown branched ester, were detected exclusively in *L. weddelli*. As behavioral evidence also suggests that tamarins can discriminate between species using olfactory cues (Epple et al., 1987), our findings suggest that such cues may depend on the presence or absence of particular compounds rather than differences in the overall composition of VOCs shared between species. However, differences in the chemical methods used across studies may limit direct comparisons, and our results should therefore be interpreted with caution. Previous analyses using headspace solid-phase microextraction in captive tamarins detected more than twice the number of our VOCs (Poirier et al., 2021a), while a study using a portable GC-MS in the same wild population detected species-specific compounds, although only a few VOCs were identified (Poirier et al., 2021b). We selected our sampling approach as advantageous for our study scope, allowing rapid collection during brief handling of wild animals without touching the gland and potentially disrupting the microbial community before sampling. However, this approach may have limited VOC detection due to low sample volumes and environmental contamination, capturing only a subset of the chemical complexity of scent gland odor profiles. Comparing methodologies and findings across studies could help clarify the advantages and limitations of different methods according to the study model and research objectives.

### ii) Multiomics did not consistently predict group membership

Contrary to our predictions, group membership was not a strong predictor for scent gland molecular profiles, and evidence differed between species. In *L. weddelli*, samples clustered according to group membership based on microbial taxonomic diversity, with pairwise comparisons showing differences in the abundance of some taxa with potential relevance for odor production (e.g., *Malassezia, Dietzia*) (Gonzalez et al., 2019; Theis et al., 2013). This could suggest that microbial taxonomic composition contributes to subtle group-level differences, although the absence of corresponding variation in predicted metabolic pathways and VOCs indicated that much of the functional potential remained conserved among groups. In *T. subgrisescens*, pairwise comparisons identified differences in the abundance of microbial genera and predicted metabolic pathways, although many of these taxa are thought to be environmentally acquired and have not been directly implicated in odor production. However, group membership explained variation in VOC composition only in *T. subgrisescens*, suggesting that group differences in odor may not be fully explained by microbial taxonomic composition alone. This contrasts with the pattern in *L. weddelli*, where group differences were limited to microbial taxonomy. Therefore, our findings mainly highlight species-specific differences in microbial contributions to scent production, rather than reflecting differences in the role of olfaction for group communication. Future studies with direct measurements of microbial- and host-derived metabolites will be essential to elucidate these mechanisms and identify the molecular features most relevant to social communication in each species.

In contrast with our findings, group-level odor signatures have been reported in primates (e.g., *Pan troglodytes* - Henkel & Setchell, 2018; *Mandrillus sphinx -* Vaglio et al., 2016; *Macaca sylvanus -* Weiß et al., 2025) and other mammals (*Lutra lutra* - Kean et al., 2017; *Suricata suricatta* - Leclaire et al., 2017; *Meles meles* - Noonan et al., 2019; *Myotis bechsteinii* - Safi & Kerth, 2003; *Crocuta* spp. - Theis et al., 2013; *Loxodonta africana* - von Dürckheim et al., 2022). In these species, group-specific odor profiles are potentially shaped by shared diet, environment, or socially mediated horizontal microbial transmission, possibly contributinf to social cohesion and/or competition over resources and territory (Brennan & Kendrick, 2006; von Dürckheim et al., 2022; Weiß et al., 2025). Some researchers have related group signatures in odor composition to genetic relatedness among group members (Boulet et al., 2009; Henkel & Setchell, 2018; Knapp et al., 2006; Leclaire et al., 2013; Porter & Moore, 1981), although evidence is contrasting (Safi & Kerth, 2003; von Dürckheim et al., 2022; Weiß et al., 2025). In our population, genetic relatedness is unlikely to closely correlate with group membership, as home ranges overlap and individuals from different groups often interact and disperse to neighbouring groups (Steinhardt, 2023; Watsa, 2013), sharing environments and potentially also contributing to horizontal transmission outside of the social group. Additionally, increasing evidence in tamarins suggests that scent may be particularly relevant to intra-group sexual communication rather than inter-group communication for territorial defence, potentially favoring cues that facilitate individual over group recognition (Brennan & Kendrick, 2006; Carboni et al., 2026b; Heymann, 1998; Miller et al., 2003; Poirier et al., 2021b). This may be particularly relevant given their cooperative breeding system with high reproductive competition (Clutton-Brock et al., 2006), and consistent with findings from other cooperative breeders, including naked mole-rats and banded mongooses, where group membership did not predict odor profiles (Jordan et al., 2010; Nichols et al., 2025). Future targeted efforts on cooperative breeders and their reliance on olfaction will be essential to determine its relative importance in intra- and inter-group social and sexual communication.

## CONCLUSIONS and FUTURE DIRECTIONS

Our study provides one of the first comprehensive molecular characterizations of scent glands in wild primates, revealing differences in microbial and olfactory patterns between closely related sympatric species. Our findings highlight two key priorities for future research: the integration of complementary multiomic datasets and a more comprehensive characterization of odor profiles. Shotgun metagenomics provides broader taxonomic coverage and a more comprehensive characterization of microbial communities than other approaches. However, this increased resolution makes direct comparisons with VOC profiles challenging, as microbial communities can comprise thousands of taxa, whereas VOC identification is constrained by lower detection and challenges in compound annotation, as described above. Future research would therefore benefit from improved reference databases and methodological approaches that enable more comprehensive and standardized characterization of both microbial and volatile profiles. Building on our integrative framework, future studies should combine behavioral observations with experimental approaches and longitudinal microbiome, metabolomic, and chemical analyses to test the causal relationships inferred from our results. In particular, integrating non-volatile metabolomics with VOC profiling would provide a more complete characterization of scent molecular profiles, contributing to clarify the role of host-microbiome interactions in shaping olfactory communication within and among groups, as well as their putative contribution to reproductive isolation.

## ACKNOWLEDGMENTS

This project was conducted with the support of the Asociación para la Conservación de la Cuenca Amazónica (ACCA) and Los Amigos Conservation Concession. We acknowledge the contributions of all field coordinators and research assistants working with Field Projects International, as well as Jhakelin Reyes Vasquez and Maria Solis Quispe for their work in supervising the Los Amigos Wildlife Conservation Laboratory. Without their contributions this research would not have been possible. We also thank Hadjira Hamou and Patricia Stroher for their help throughout the project with advice on wet lab protocols, analysis, and research permits.

## FUNDERS

The study received funding from: the Natural Sciences and Engineering Research Council of Canada (RGPIN-2017-03782 to ADM), the Canada Foundation for Innovation and Canada Research Chairs program (950-231257 to ADM), the Explorers Club (Exploration Fund Grant, to SC), the International Primatological Society (Research Grant, to SC), SigmaXi (Grant in Aid of Research G20240315-8618 to SC), the Killam Trust (Killam Doctoral Scholarship to SC), Field Projects International, and the Leakey Foundation (Research Grant 10037034 to ACP and ADM).

### Ethical statement

The study followed the guidelines of the Canadian Council of Animal Care and was approved by the Animal Care Committees of the University of Calgary (ACC19-0167) and of Washington University (#21-0084). The annual mark-recapture program and the sample collection conducted by FPI were authorized by the Peruvian Servicio Nacional Forestal y de Fauna Silvestre (SERFOR-D000116-2021 and SERFOR-D000094-2022). Importation of samples to Canada was approved by the Canadian Food Inspection Agency (A-2022-05138-1; A-2024-00080-1).

### Data Accessibility Statement

Raw sequence reads are deposited in the NCBI SRA (PRJNA1260568 and PRJNA1536639). Odor sample metadata are publicly available at https://github.com/AlicePoirier/Carboni-et-al_Microbial-contributions-to-species-specific-olfactory-communication-in-wild-tamarins. Sample metadata are stored in the same BioProject and in Supplementary Material 1.

### Benefit-Sharing Statement

Benefits generated: This research fostered international collaboration by partnering with the non-profit organizations Field Projects International and the In Situ Lab initiative. These partnerships enabled the collection of samples through coordinated efforts among researchers from Canada, the United States, and Peru. All collaborators are acknowledged as co-authors on resulting publications, and the data generated have been made publicly available through the database described above.

### Author Contributions

S. Carboni contributed to research design, funding acquisition, data collection, data curation, investigation, data analysis, and writing the original draft. A.C. Poirier contributed to research design, funding acquisition, data collection, investigation, data analysis, and data curation. M. Watsa and G. Erkenswick contributed to funding acquisition, project administration, and investigation. A.D. Melin contributed to funding acquisition, research design, supervision, and major editing of the manuscript. All authors reviewed and approved the final version of the manuscript for submission.

