## Supplementary material for "Multiomic evidence for microbial contributions to species-specific olfactory communication in wild tamarins": Suppl. Materials 2: SupplementaryMaterials2_GlandMultiomic.docx

**SUPPLEMENTARY MATERIALS 2**

**Suppl. Table 1.** Description of thermal desorption–gas chromatography-mass spectrometry (TD–GC-MS) analyses and chemical data processing of tamarin scent gland odor collected via probe sampling, and example of a resulting chemical profile.

| **TD– CG-MS methods** | - GC-MS system: Shimadzu QP2020 NX GC-MS linked to a TD-30R thermal desorption unit (TDU) - Helium used as carrier gas at a flow rate of 1 mL/min - Probe samples placed into clean TD tubes for analysis (unpacked stainless steel TD tube, 1/4 in. × 3 1/2 in., Supelco, Bellefont, USA) - TDU parameters: tube desorption at 300°C in splitless mode, held for 15 min - Injector parameters: sample injection at 250°C, held for 6 min - GC system: DB-5 low-polar capillary column (30 m × 0.25 mm × 0.25 µm, Agilent) - GC parameters: Initial oven temperature set to 40°C, held for 1 min and then increased by 7°C/min to 300°C, held for 8 min (total run time = 38 min) - MS system: electron ionization (scan range 30-500 Da) - MS parameters: transfer line temperature set to 250°C and ion source temperature to 230°C; detection window set to 1–38 min |
| --- | --- |
| **Example chemical profiles** | 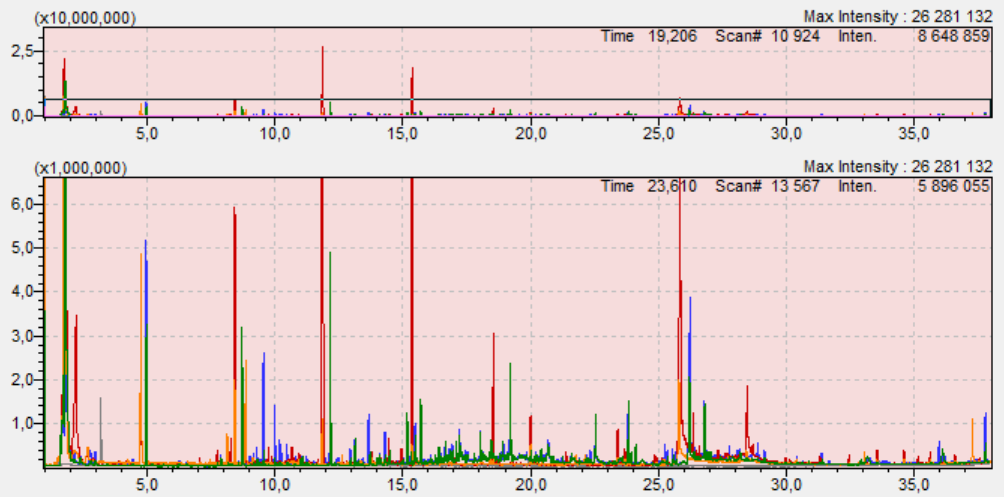  *(green)* adult female saddleback tamarin (group S7)  *(blue)* adult female emperor tamarin (group IC)  *(orange)* adult male emperor tamarin (group MI)  *(red)* air control  *(grey)* instrument blank (i.e., empty run to control for carry over in colum) |
| **Chemical data processing** | - Automatic peak detection, deconvolution and integration in AMDIS 2.73 with the following parameters: high resolution, sensitivity and shape requirement, component width = 12, single adjacent peak - Manual correction of peak assignment by considering mass-to-charge (m/z) ratios and retention times of peaks - Creation of a custom peak library applied to all profiles in GC-MS Solutions v.4.20 (Shimadzu, Kyoto, Japan) to extract the peak areas of all specified peaks within their assigned RT range (i.e. within ± 50% of the library range entry for each peak) and a signal-to-noise ratio threshold ≥1 - Manual peak selection steps in R v.4.5.2:   - Exclusion of all peaks found in ≥ 20% of blank or control profiles   - Exclusion of all peaks found in higher amounts in blanks and controls   - Exclusion of rare peaks, i.e. found in < 3 samples   - Exclusion of samples with < 2 peaks - Tentative VOC identification using the NIST 14 mass spectral library on the basis of their mass spectra and retention indices calculated based on an n-alkane reference mixture analysed under identical conditions (NIST identity match threshold = 80%) |

**Suppl. Table 2.** Volatile organic compounds (VOCs) retrieved from the suprapubic gland of wild emperor and Weddell’s saddleback tamarins, their tentative identity, retention time ± standard deviation (RT ± sd, in min), chemical class, and references to these compounds in the mammalian literature.

| **VOC #** | **Tentative identity** | **RT ± sd** | **Chemical class** | **Mention in the mammalian literature (mainly in glandular secretions and body odor)** |
| --- | --- | --- | --- | --- |
| 1 | Methyl propanoate | 2.447 ± 0.037 | Methyl ester | In faeces of house mice, *Mus musculus* (1) |
| 2 | Unidentified compound 01 | 2.916 ± 0.049 | Unk. | N/A |
| 3 | Acetic acid | 3.186 ± 0.027 | Fatty acid (carboxylic acid) | In vaginal secretions of female bonobos, *Pan paniscus* (2), and rhesus macaques, *Macaca mulatta* (3); in scent-marks of common marmosets, *Callithrix jacchus* (4,5) and giant pandas, *Ailuropoda melanoleuca* (6); in anogenital gland secretion of Milne-Edwards' sifaka, *Propithecus edwarsi* (7); in anal gland secretions of small Indian mongooses, *Herpestes javanicus auropunctatus* (8) and diverse canids (9); in sternal gland secretions of male mandrills, *Mandrillus sphinx* (10), and koalas, *Phascolarctos cinereus* (11); in interdigital gland secretions of male white-tailed deer, *Odocoileus virginianus* (12); in body odor of meerkats, *Suricata suricatta* (13), and humans, *Homo sapiens* (14,15) |
| 4 | Methylbenzene | 3.995 ± 0.063 | Aromatic hydrocarbon | In vaginal secretions of female olive baboons, *Papio anubis* (16); in scent-marks of giant pandas (6); in brachial gland secretions of ring-tailed lemurs, *Lemur catta* (17); in anal gland secretions of European badgers, *Meles meles* (18); in tail gland secretions of red foxes, *Vulpes vulpes* (19); in body odor of common marmosets (20), ring-tailed lemurs (17), humans (21) and waterbuck, *Kobus defassa* (22) |
| 5 | Butanoic acid | 4.597 ± 0.067 | Fatty acid | In vaginal secretions of female bonobos (2), olive baboons (16), and rhesus macaques (3); in scent-marks of common marmosets (4) and giant pandas (6); in anal gland secretions of canids (9), badgers (18) and mongooses (8,23,24); in cheek gland secretions of dwarf hamster, *Phodopus sungorus sungorus* (25); in body odor of humans (15) and mice (26) |
| 6 | Unidentified compound 02 | 5.131 ± 0.030 | Unk. | N/A |
| 7 | Unidentified compound 03 | 5.224 ± 0.009 | Unk. | N/A |
| 8 | Unidentified compound 04 | 6.849 ± 0.026 | Unk. | N/A |
| 9 | Unidentified compound 05 | 7.014 ± 0.056 | Unk. | N/A |
| 10 | Unk. benzene derivative 1 | 7.377 ± 0.006 | Benzene derivative | N/A |
| 11 | Benzaldehyde | 7.797 ± 0.022 | Aromatic aldehyde | In vaginal secretions of female olive baboons (16); in scent-marks of tamarins (27), common marmosets (4,5), red-ruffed lemurs, *Varecia variegata rubra* (28), white-faced saki monkeys, *Pithecia pithecia* (29) and giant pandas (6); in suprapubic gland secretions of silvery marmosets, *Mico argentatus* (29); in subcaudal gland secretion of owl monkeys, *Aotus* spp. (30); in anal gland secretions of tamarins (27), canids (9), badgers (18), wolverines, *Gulo gulo* (31), ferrets, *Mustela furo* (32) and crested porcupines, *Hystrix cristata* (33); in sternal gland secretions of male mandrills (10); in chin gland secretions of European rabbit, *Oryctolagus cuniculus* (34); in interdigital gland secretions of male white-tailed deer (12); in cheek and forehead odor of felids (35); in body odor of common marmosets (20), humans (15,21,36,37,38), mice (26) and reticulated giraffes, *Giraffa camelopardalis reticulata* (39) |
| 12 | Benzonitrile | 8.285 ± 0.018 | Aromatic nitrile | In faeces of Eurasian otters, *Lutra lutra* (40) |
| 13 | D-Limonene | 9.131 ± 0.038 | Monoterpene | In vaginal secretions of female olive baboons (16); in sternal gland secretions of mandrills (41); in body odor of humans (37) and common marmosets (42) |
| 14 | Benzeneacetaldehyde | 9.723 ± 0.064 | Aromatic aldehyde | In suprapubic gland secretions of tamarins and marmosets (27,29); in scent-marks of tamarins (29) and giant pandas (6) |
| 15 | Unk. C11 branched alkane | 9.759 ± 0.019 | Alkane | N/A |
| 16 | Unidentified compound 06 | 10.032 ± 0.113 | Unk. | N/A |
| 17 | 1-Methyl-2-(3-methylpentyl)cyclopropane | 10.494 ± 0.034 | Methyl cycloalkane | None reported |
| 18 | Unidentified compound 07 | 10.761 ± 0.066 | Unk. | N/A |
| 19 | Unk. C12 branched alkane 1 | 10.815 ± 0.075 | Alkane | N/A |
| 20 | Unidentified compound 08 | 10.999 ± 0.020 | Unk. | N/A |
| 21 | Nonanal | 11.126 ± 0.057 | Aldehyde | In vaginal secretions of female white-tailed deer (43); in scent-marks of common marmosets (5) and giant pandas (6); in anal gland secretions of ferrets (32) and crested porcupines (33); in tail gland secretions of red foxes (19); in brachial gland secretions of ring-tailed lemurs (17); in sternal gland secretions of male mandrills (10,41) and koalas (11); in cheek and forehead odor of felids (35); in interdigital gland secretions of male white-tailed deer (12) and red hartebeest, *Alcelaphus buselaphus caama* (44); in metatarsal gland secretions of sika deer, *Cervus nippon* (45), and tarsal gland secretions of reindeer, *Rangifer tarandus* (46); in preorbital gland secretions of sunis, *Neotragus moschatus* (47); in body odor of common marmosets (20), humans (14,15,36,37,38,48), meerkats (13), African buffalos, *Syncerus caffer* (22) and reticulated giraffes (39) |
| 22 | Unidentified compound 09 | 11.316 ± 0.023 | Unk. | N/A |
| 23 | Unk. C12 branched alkane 2 | 11.324 ± 0.006 | Alkane | N/A |
| 24 | Unidentified compound 10 | 11.502 ± 0.052 | Unk. | N/A |
| 25 | 2,2-Dimethyldecane | 11.558 ± 0.061 | Methyl alkane | In human breath (49) |
| 26 | Unidentified compound 11 | 11.929 ± 0.053 | Unk. | N/A |
| 27 | Octanoic acid | 12.378 ± 0.060 | Fatty acid | In scent-marks of giant pandas (6); in anal gland secretions of black-backed jackals, *Canis mesomelas* (9), wolverines (31), meerkats (50), and crested porcupines (33); in tail gland secretions of red foxes (19); in sternal gland secretions of male koalas (11); in cheek and forehead odor of felids (35); in dorsal gland secretions of white-lipped peccaries, *Tayassu pecari* (51); in pedal gland secretions of brown bear, *Ursus arctos* (52); in interdigital gland secretions of male white-tailed deer (12); in metatarsal gland secretions of sika deer (45); in preorbital gland secretions of sunis (47); in body odor of humans (14,36,48), mice (53), and waterbucks, *Kobus defassa* (22) |
| 28 | Unk. benzene derivative 2 | 12.396 ± 0.043 | Benzene derivative | N/A |
| 29 | Unidentified compound 12 | 12.669 ± 0.015 | Unk. | N/A |
| 30 | Nonylcyclopropane | 12.944 ± 0.013 | Cycloalkane | None reported |
| 31 | 1,4:3,6-Dianhydro-α-D-glucopyranose | 13.077 ± 0.083 | Bicyclic carbohydrate derivative | None reported |
| 32 | 3,5-Dimethylundecane | 13.489 ± 0.104 | Methyl alkane | 3,6-Dimethylundecane in mandrill sternal gland secretions (41); 3,7-dimethylundecane in human breath (54) |
| 33 | Methenamide | 13.603 ± 0.021 | Heterocyclic amide | None reported |
| 34 | 4,6-Dimethyldodecane | 13.905 ± 0.051 | Methyl alkane | In mandrill sternal gland secretions (41) |
| 35 | Unidentified compound 13 | 16.691 ± 0.026 | Unk. | N/A |
| 36 | Unk. C14 branched alkane | 16.757 ± 0.049 | Alkane | N/A |
| 37 | Unidentified compound 14 | 16.950 ± 0.033 | Unk. | N/A |
| 38 | 2-Methyldodecan-2-ol | 17.672 ± 0.019 | Methyl alcohol | None reported |
| 39 | 2,2-Dimethyltetradecane | 19.194 ± 0.034 | Methyl alkane | None reported |
| 40 | Unidentified compound 15 | 19.386 ± 0.013 | Unk. | N/A |
| 41 | Tridecan-1-ol | 20.201 ± 0.058 | Alcohol | In mandrill sternal gland secretions (10); in anal gland secretions of mongooses (23); in metatarsal gland secretions of sika deer (45); in human body odor (55) |
| 42 | Unidentified compound 16 | 20.766 ± 0.003 | Unk. | N/A |
| 43 | Unk. benzene derivative 3 | 21.074 ± 0.032 | Benzene derivative | N/A |
| 44 | Unk. branched alcohol | 21.180 ± 0.017 | Alcohol | N/A |
| 45 | Unidentified compound 17 | 22.081 ± 0.021 | Unk. | N/A |
| 46 | Unidentified compound 18 | 22.175 ± 0.045 | Unk. | N/A |
| 47 | Unidentified compound 19 | 23.098 ± 0.029 | Unk. | N/A |
| 48 | Pentanedecanitrile | 23.836 ± 0.042 | Nitrile | None reported |
| 49 | Pentadecanoic acid | 23.859 ± 0.015 | Fatty acid | In labial and scrotal gland secretions of ring-tailed lemurs (56); in spur and cloacal wax secretions of short-beaked echidnas, *Tachyglossus aculeatus* (57); in giant panda anogenital gland secretions (58,59); in anal gland secretions of canids (9), ferrets (32), meerkats (50), wolverines (31), crested porcupines (33), Eurasian beavers, *Castor fiber* (60); in tail gland secretions of red foxes (19); in dorsal gland secretions of white-lipped peccaries (51); in interdigital gland secretions of male red hartebeest (44); in metatarsal gland secretions of sika deer (45); in occipital gland secretions of bactrian camels, *Camelus bactrianus* (61); in human body odor (14,15) |
| 50 | (E)-Hexadec-9-en-1-ol | 24.828 ± 0.030 | Fatty alcohol | In mice preputial gland secretions (62) |
| 51 | Unk. branched ester | 25.490 ± 0.046 | Ester | N/A |
| 52 | Unidentified compound 20 | 25.972 ± 0.040 | Unk. | N/A |
| 53 | Unk. branched carboxylic acid | 26.472 ± 0.012 | Fatty acid | N/A |
| 54 | Nonanamide | 27.585 ± 0.050 | Fatty acid amide | In urine of bobcats, *Lynx rufus* (63) |
| 55 | Unidentified compound 21 | 27.953 ± 0.022 | Unk. | N/A |
| 56 | Unidentified compound 22 | 28.689 ± 0.020 | Unk. | N/A |
| 57 | Unk. amide 1 | 28.975 ± 0.003 | Amide | N/A |
| 58 | Unk. amide 2 | 30.230 ± 0.028 | Amide | N/A |
| 59 | Unidentified compound 23 | 30.761 ± 0.091 | Unk. | N/A |
| **References cited in table:**   1. Reade, S. P. (2016). *Volatile organic compound profiling of mouse models of intestinal inflammation and colon carcinogenesis* [PhD thesis]. University of Liverpool. 2. Matsumoto-Oda, A., et al. (2003). Vaginal fatty acids produced by chimpanzees during menstrual cycles. *Folia Primatologica*, *74*(2), 75–79.<https://doi.org/10.1159/000070000> 3. Michael, R. P., Keverne, E. B., & Bonsall, R. W. (1971). Pheromones: Isolation of male sex attractants from a female primate. *Science*, *172*, 964–966. 4. Smith, T. E. (2001). Female marmoset monkeys (*Callithrix jacchus*) can be identified from the chemical composition of their scent marks. *Chemical Senses*, *26*(5), 449–458.<https://doi.org/10.1093/chemse/26.5.449> 5. Thompson, C. L., et al. (2020). What smells? Developing in‐field methods to characterize the chemical composition of wild mammalian scent cues. *Ecology and Evolution*, *10*(11), 4691–4701.<https://doi.org/10.1002/ece3.6224> 6. Hagey, L., & Macdonald, E. (2003). Chemical cues identify gender and individuality in giant pandas (*Ailuropoda melanoleuca*). *Journal of Chemical Ecology*, 10.<https://doi.org/10.1023/A:1024225806263> 7. Morelli, T. L., et al. (2013). Relatedness communicated in lemur scent. *Naturwissenschaften*, *100*(8), 769–777.<https://doi.org/10.1007/s00114-013-1074-x> 8. Miyazaki, T., et al. (2018). Identification of 2-phenylethanol with a rose-like odor from anal sac secretions of the small Indian mongoose (*Herpestes auropunctatus*). *Bioscience, Biotechnology, and Biochemistry*, *82*(2), 232–237.<https://doi.org/10.1080/09168451.2017.1419854> 9. Apps, P., Mmualefe, L., & McNutt, J. W. (2012). Identification of volatiles from the secretions and excretions of African wild dogs (*Lycaon pictus*). *Journal of Chemical Ecology*, *38*(11), 1450–1461.<https://doi.org/10.1007/s10886-012-0206-7> 10. Vaglio, S., et al. (2016). Sternal gland scent-marking signals sex, age, rank, and group identity in captive mandrills. *Chemical Senses*, bjv077.<https://doi.org/10.1093/chemse/bjv077> 11. Salamon, M., & Davies, N. W. (1998). Identification and variation of volatile compounds in sternal gland secretions of male koalas (*Phascolarctos cinereus*). *Journal of Chemical Ecology*, *24*(10), 1659–1676.<https://doi.org/10.1023/A:1020868528762> 12. Gassett, J. W., et al. (1996). Volatile compounds from interdigital gland of male white-tailed deer (*Odocoileus virginianus*). *Journal of Chemical Ecology*, *22*(9), 1689–1696.<https://doi.org/10.1007/BF02272407> 13. Weiß, B. M., et al. (2018). A non‐invasive method for sampling the body odour of mammals. *Methods in Ecology and Evolution*, *9*(2), 420–429.<https://doi.org/10.1111/2041-210X.12888> 14. Akin, J. (2005). *Characterization of human skin emanations by solid phase microextraction (SPME) extraction of volatiles and subsequent analysis by gas chromatography-mass spectrometry (GC-MS)* [BSc thesis]. Massachusetts Institute of Technology. 15. Gallagher, M., et al. (2008). Analyses of volatile organic compounds from human skin. *British Journal of Dermatology*, *159*(4), 780–791.<https://doi.org/10.1111/j.1365-2133.2008.08748.x> 16. Vaglio, S., et al. (2021). Olfactory signals and fertility in olive baboons. *Scientific Reports*, *11*(1), 8506.<https://doi.org/10.1038/s41598-021-87893-6> 17. Knapp, L. A., Robson, J., & Waterhouse, J. S. (2006). Olfactory signals and the MHC: A review and a case study in *Lemur catta*. *American Journal of Primatology*, *68*(6), 568–584.<https://doi.org/10.1002/ajp.20253> 18. Noonan, M. J., et al. (2019). Knowing me, knowing you: Anal gland secretion of European badgers (*Meles meles*) codes for individuality, sex and social group membership. *Journal of Chemical Ecology*, *45*(10), 823–837.<https://doi.org/10.1007/s10886-019-01113-0> 19. McLean, S., Davies, N. W., & Nichols, D. S. (2019). Scent chemicals of the tail gland of the red fox, *Vulpes vulpes*. *Chemical Senses*, *44*(3), 215–224.<https://doi.org/10.1093/chemse/bjz009> 20. Kücklich, M., Möller, M., Marcillo, A., Einspanier, A., Weiß, B. M., Birkemeyer, C., & Widdig, A. (2017). Different methods for volatile sampling in mammals. *PLOS ONE*, *12*(8), e0183440.<https://doi.org/10.1371/journal.pone.0183440> 21. Curran, A. M., Rabin, S. I., & Furton, K. G. (2005). Analysis of the uniqueness and persistence of human scent. *Forensic Science Communications*, *7*(2), 1–8. 22. Gikonyo N. K. et al. (2002). Odor composition of preferred (buffalo and ox) and nonpreferred (waterbuck) hosts of some savanna tsetse flies. *Journal of Chemical Ecology*. 25(5). <https://doi.org/10.1023/a:1015205716921> 23. Gorman, M. L. (1976). A mechanism for individual recognition by odour in *Herpestes auropunctatus* (Carnivora: Viverridae). *Animal Behaviour*, *24*(1), 141–145.<https://doi.org/10.1016/S0003-3472(76)80107-8> 24. Jordan, N. R., et al. (2010). Scent marking within and between groups of wild banded mongooses. *Journal of Zoology*, *280*(1), 72–83.<https://doi.org/10.1111/j.1469-7998.2009.00646.x> 25. Burger, B. V., et al. (2001). Mammalian exocrine secretions XVI. Constituents of secretion of supplementary sacculi of dwarf hamster, *Phodopus sungorus sungorus*. *Journal of Chemical Ecology*, 27(6). <https://doi.org/10.1023/a:1010380315961> 26. Röck, F., et al. (2006). Comparative analysis of volatile constituents from mice and their urine. *Journal of Chemical Ecology*, *32*(6), 1333–1346.<https://doi.org/10.1007/s10886-006-9091-2> 27. Poirier, A. C., et al. (2021). Scent marks signal species, sex, and reproductive status in tamarins (*Saguinus* spp., Neotropical primates). *Chemical Senses*, *46*, 1–12.<https://doi.org/10.1093/chemse/bjab008> 28. Janda, E. D., et al. (2019). Sex differences in scent-marking in captive red-ruffed lemurs. *American Journal of Primatology*, *81*(1), e22951.<https://doi.org/10.1002/ajp.22951> 29. Poirier, A. C. (2019). *Scent-marking behaviour and semiochemistry in the Callitrichidae* [PhD thesis]. Anglia Ruskin University, UK. 30. Spence-Aizenberg, A. (2017). *Olfactory communication, mate choice, and reproduction in a pair-bonded primate (*Aotus *spp.)* [PhD thesis]. University of Pennsylvania. 31. Wood, W. F., Terwilliger, M. N., & Copeland, J. P. (2005). Volatile compounds from anal glands of the wolverine, *Gulo gulo*. *Journal of Chemical Ecology*, *31*(9), 2111–2117.<https://doi.org/10.1007/s10886-005-6080-9> 32. Zhang, J. X., et al. (2005). Putative chemosignals of the ferret (*Mustela furo*) associated with individual and gender recognition. *Chemical Senses*, *30*(9), 727–737.<https://doi.org/10.1093/chemse/bji065> 33. Massolo, A., Dani, F. R., & Bella, N. (2009). Sexual and individual cues in the peri-anal gland secretum of crested porcupines (*Hystrix cristata*). *Mammalian Biology*, *74*(6), 488–496.<https://doi.org/10.1016/j.mambio.2009.07.004> 34. Hayes, R. A., Richardson, B.J ., Wyllie, S. G. (2002). Semiochemicals and social signaling in the wild European rabbit in Australia: I. Scent profiles of chin gland secretion from the field. *Journal of Chemical Ecology*, 28(2):363–384. <https://doi.org/10.1023/a:1017942426467> 35. Soini, H. A., et al. (2012). Investigation of scents on cheeks and foreheads of large felines in connection to the facial marking behavior. *Journal of Chemical Ecology*, *38*(2), 145–156.<https://doi.org/10.1007/s10886-012-0075-0> 36. Logan, J. G., et al. (2008). Identification of human-derived volatile chemicals that interfere with attraction of *Aedes aegypti* mosquitoes. *Journal of Chemical Ecology*, *34*(3), 308–322.<https://doi.org/10.1007/s10886-008-9436-0> 37. Medrano, A. C., et al. (2024). Chemical characterization of human body odor headspace components. *Separations*, *11*(3), 85.<https://doi.org/10.3390/separations11030085> 38. Mochalski, P., et al. (2012). Temporal profiling of human urine VOCs and its potential role under the ruins of collapsed buildings. *Toxicology Mechanisms and Methods*, *22*(7), 502–511.<https://doi.org/10.3109/15376516.2012.682664> 39. Wood, W. F., & Weldon, P. J. (2002). The scent of the reticulated giraffe (*Giraffa camelopardalis reticulata*). *Biochemical Systematics and Ecology*, *30*(10), 913–917.<https://doi.org/10.1016/S0305-1978(02)00037-6> 40. Kean, E. F., Chadwick, E. A., & Müller, C. T. (2015). Scent signals individual identity and country of origin in otters. *Mammalian Biology*, *80*(2), 99–105.<https://doi.org/10.1016/j.mambio.2014.12.004> 41. Setchell, J. M., et al. (2010). Chemical composition of scent-gland secretions in an Old World monkey (*Mandrillus sphinx*): Influence of sex, male status, and individual identity. *Chemical Senses*, *35*(3), 205–220.<https://doi.org/10.1093/chemse/bjp105> 42. Kücklich M. et al. (2019). Chemical cues of female fertility states in a non-human primate. *Scientific Reports*, 9(1):13716. <https://doi.org/10.1038/s41598-019-50063-w> 43. Jemiolo, B., et al. (1995). Putative chemical signals from white-tailed deer (*Odocoileus virginianus*). Urinary and vaginal mucus volatiles excreted by females during breeding season. *Journal of Chemical Ecology*, *21*(6), 869–879.<https://doi.org/10.1007/BF02033467> 44. Reiter, B., Burger, B. V., & Dry, J. (2003). Mammalian exocrine secretions. XVIII: Chemical characterization of interdigital secretion of red hartebeest, *Alcelaphus buselaphus caama*. *Journal of Chemical Ecology*, 29(10):2235–52. <https://doi.org/10.1023/a:1026218213151> 45. Wood, W. F. (2003). Volatile components in metatarsal glands of Sika deer, *Cervus nippon*. *Journal of Chemical Ecology*, *29*(12), 2729–2733.<https://doi.org/10.1023/B:JOEC.0000008016.44795.8a> 46. Andersson, G., et al. (1975). Volatile compounds from the tarsal scent gland of reindeer (*Rangifer tarandus*). *Journal of chemical ecology*, *1*(2), 275–281.<https://doi.org/10.1007/bf00987876> 47. Stander, M. A., Burger, B. V., & Roux, M. L. (2001). Mammalian Exocrine Secretions. XVII: Chemical Characterization of Preorbital Secretion of Male Suni, *Neotragus moschatus*. *Journal of Chemical Ecology*, 28:89–101. <https://doi.org/10.1023/A:1013562818965> 48. Owsienko, D., et al. (2024). Body odor samples from infants and post-pubertal children differ in their volatile profiles. *Communications Chemistry*, *7*(1), 1–10.<https://doi.org/10.1038/s42004-024-01131-4> 49. Peng G. et al. (2010). Detection of lung, breast, colorectal, and prostate cancers from exhaled breath using a single array of nanosensors. *British Journal of Cancer*, 103(4):542–551. <https://doi.org/10.1038/sj.bjc.6605810> 50. Leclaire, S., et al. (2017). Social odours covary with bacterial community in the anal secretions of wild meerkats. *Scientific Reports*, *7*(1), 3240.<https://doi.org/10.1038/s41598-017-03356-x> 51. Waterhouse, J. S., et al. (2001). Volatile components in dorsal gland secretions of the white-lipped peccary, *Tayassu pecari*, from Bolivia. *Journal of Chemical Ecology*, 27:2459–2469. <https://doi.org/10.1023/A:1013675431162> 52. Sergiel, A., et al. (2017). Histological, chemical and behavioural evidence of pedal communication in brown bears. *Scientific Reports*, *7*(1), 1052.<https://doi.org/10.1038/s41598-017-01136-1> 53. Brechbühl, J., et al. (2013). Mouse alarm pheromone shares structural similarity with predator scents.  *Proceedings of the National Academy of Sciences*, 110(12):4762–4767. <https://doi.org/10.1073/pnas.1214249110> 54. Dragonieri, S., et al. (2007). An electronic nose in the discrimination of patients with asthma and controls. *Journal of Allergy and Clinical Immunology*. 120(4):856–862. <https://doi.org/10.1016/j.jaci.2007.05.043> 55. Penn, D. J., et al. (2007). Individual and gender fingerprints in human body odour. *Journal of The Royal Society Interface*, *4*(13), 331–340.<https://doi.org/10.1098/rsif.2006.0182> 56. Scordato, E. S., Dubay, G., & Drea, C. M. (2007). Chemical composition of scent marks in the ringtailed Lemur (*Lemur catta*): Glandular differences, seasonal variation, and individual signatures. *Chemical Senses*, *32*(5), 493–504.<https://doi.org/10.1093/chemse/bjm018> 57. Harris, R. L., et al. (2014). Chemical signals in the echidna: Differences between seasons, sexes, individuals and gland types. *Journal of Zoology*, *293*(3), 171–180.<https://doi.org/10.1111/jzo.12133> 58. Yuan, H., et al. (2004). Anogenital gland secretions code for sex and age in the giant panda, *Ailuropoda melanoleuca*. *Canadian Journal of Zoology*, *82*(10), 1596–1604.<https://doi.org/10.1139/z04-144> 59. Zhang, J.-X., et al. (2008). Potential chemosignals in the anogenital gland secretion of giant pandas, *Ailuropoda melanoleuca*, associated with sex and individual identity. *Journal of Chemical Ecology*, *34*(3), 398–407.<https://doi.org/10.1007/s10886-008-9441-3> 60. Rosell, F., & Sundsdal, L. J. (2001). Odorant source used in Eurasian beaver territory marking. *Journal of Chemical Ecology*, 27:2471–2491. <https://doi.org/10.1023/A:1013627515232> 61. Ayorinde, F., et al. (1982). Volatile components of the occipital gland secretion of the bactrian camel (Camelus bactrianus). *Journal of Chemical Ecology*, 8(1):177–183. <https://doi.org/10.1007/BF00984014> 62. Zhang, J.-X., et al. (2007). Putative chemical signals about sex, individuality, and genetic background in the preputial gland and urine of the house mouse (*Mus musculus*). *Chemical Senses*, *32*(3), 293–303.<https://doi.org/10.1093/chemse/bjl058> 63. Mattina, M. J. I., Pignatello, J. J., & Swihart, R. K. (1991). Identification of volatile components of bobcat (*Lynx rufus*) urine. *Journal of Chemical Ecology*, *17*(2), 451–462.<https://doi.org/10.1007/BF00994344> | | | | |

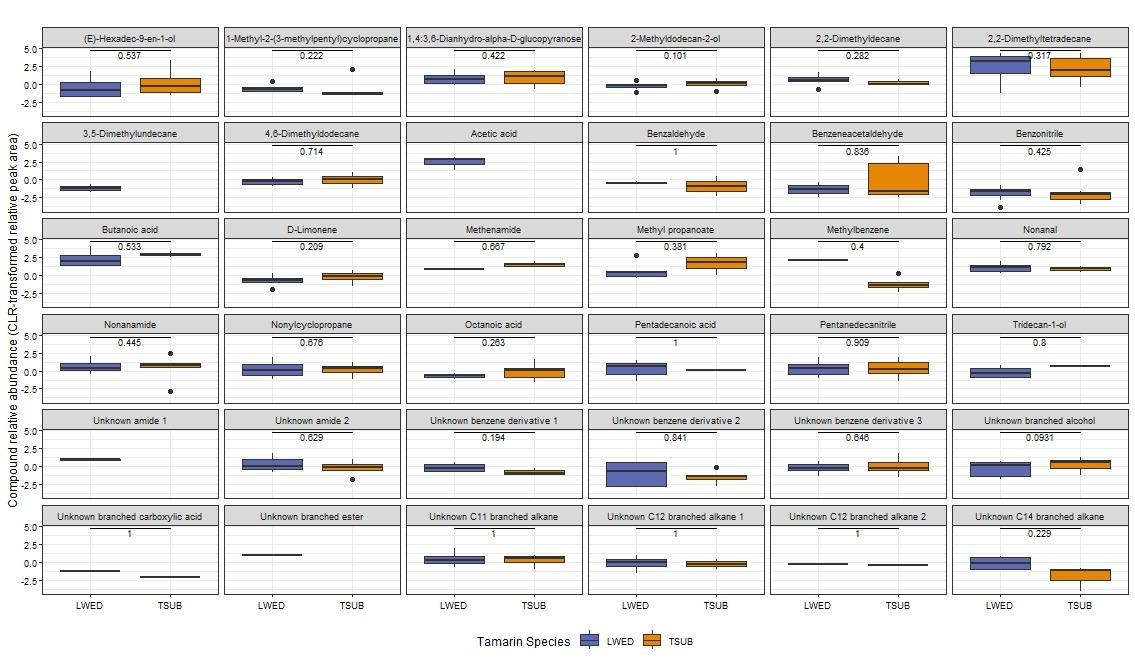

**Suppl. Figure 1.** Relative abundances of all identified VOCs, normalized by CLR transformation of relative peak area, and separated by tamarin species. For compounds detected in both species, Wilcoxon rank-sum tests were performed, with Holm-adjusted p-values shown above the boxplots. None of the comparisons were significant.
