## Supplementary material for "Multiomic evidence for microbial contributions to species-specific olfactory communication in wild tamarins": Suppl. Materials 3: SupplementaryMaterials3_GlandMultiomic.docx

**SUPPLEMENTARY MATERIAL 3**

**Suppl. Table 1.** Results from GLMMs predicting two measures of alpha diversity calculated from microbial taxonomy of 135 samples:

- Shannon Model: *Shannon index ~ Tamarin species + Sequencing year + Year of data collection + Read count per sample + (1 | Animal identity).*

Data distribution: Gamma with a log-link function.

- Richness Model: *Richness ~ Tamarin species + Sequencing year + Year of data collection + Read count per sample + (1 | Animal identity).*

Data distribution: Poisson with a log-link function.

Significance: p<0.05 (*), p<0.01 (**), p<0.001 (***). The AICc values for each alternative model and their respective null models are reported at the bottom of the table; in both cases, the alternative models outperformed the null models.

| **Shannon Model: Shannon diversity of microbial taxonomy as the response variable** | | | | | | |
| --- | --- | --- | --- | --- | --- | --- |
| **Explanatory variable** | **Estimate** | **SE** | ***t-value*** | ***p-value*** | **Random effect: Animal ID** | **R² values** |
| Intercept | 1.941 | 0.023 | 85.015 | < 2e-16* | σ^2^: 0.0041 | R²m: 0.237 |
| Species: TSUB^a^ | 0.083 | 0.025 | 3.300 | < 0.001* | sd: 0.064 | R²c: 0.513 |
| Sequencing Run: 2025^b^ | -0.105 | 0.037 | -2.794 | 0.0052* |  |  |
| Year: 2023^c^ | 0.041 | 0.036 | 1.147 | 0.251 |  |  |
| Year: 2024^c^ | 0.050 | 0.036 | 1.373 | 0.170 |  |  |
| Read Count (scaled) | -0.024 | 0.010 | -2.381 | 0.017* |  |  |
| **Richness Model: Richness of microbial taxonomy as the response variable** | | | | | | |
| **Explanatory variable** | **Estimate** | **SE** | ***t-value*** | ***p-value*** | **Random effect: Animal ID** | **R² values** |
| Intercept | 8.876 | 0.004 | 2330.98 | < 2e-16* | σ^2^: 0.0003 | R²m: 0.309 |
| Species: TSUB^a^ | 0.006 | 0.004 | 1.305 | 0.192 | sd: 0.019 | R²c: 0.801 |
| Sequencing Run: 2025^b^ | -0.008 | 0.003 | -2.594 | 0.009* |  |  |
| Read Count (scaled) | 0.013 | 0.002 | 8.172 | 3e-06* |  |  |

Reference category:  ^a^ LWED; ^b^ 2023; ^c^ 2022

**AICc values for Model Shannon:**

Null model: 307.9

Alternative model: 293.0

**AICc values for Model Richness:**

Null model: 2133.9

Alternative model: 2050.8

**Suppl. Table 2.** Results from GLMMs predicting two measures of alpha diversity calculated from microbial metabolic pathways of 135 samples:

- Shannon Model: *Shannon index ~ Tamarin species + Sequencing year + Year of data collection + (1 | Animal identity).*

Data distribution: Gamma with a log-link function.

- Richness Model: *Richness ~ Tamarin species + Sequencing year + Year of data collection + (1 | Animal identity).*

Data distribution: Poisson with a log-link function.

Significance: p<0.05 (*), p<0.01 (**), p<0.001 (***). The AICc values for each alternative model and their respective null models are reported at the bottom of the table. For the Shannon model, the null model provided a better fit than the alternative model, whereas for the Richness model, the alternative model outperformed the null model.

| **Shannon Model: Shannon diversity of pathways abundance as the response variable** | | | | | | |
| --- | --- | --- | --- | --- | --- | --- |
| **Explanatory variable** | **Estimate** | **SE** | ***t-value*** | ***p-value*** | **Random effect: Animal ID** | **R² values** |
| Intercept | 1.689 | 0.009 | 188.710 | < 2e-16* | σ^2^: 0.0007  sd: 0.027 | R²m: 0.047 |
| Species: TSUB^a^ | -0.004 | 0.010 | -0.380 | 0.704 |  | R²c: 0.412 |
| Sequencing Run: 2025^b^ | -0.029 | 0.014 | -1.983 | 0.047* |  |  |
| Year: 2023^c^ | 0.008 | 0.014 | 0.602 | 0.547 |  |  |
| Year: 2024^c^ | 0.016 | 0.014 | 1.105 | 0.269 |  |  |
| **Richness Model: Richness of pathways abundance as the response variable** | | | | | | |
| **Explanatory variable** | **Estimate** | **SE** | ***t-value*** | ***p-value*** | **Random effect: Animal ID** | **R² values** |
| Intercept | 5.913 | 0.036 | 163.954 | < 2e-16* | σ^2^: 0.057  sd: 0.240 | R²m: 0.086 |
| Species: TSUB^a^ | -0.064 | 0.050 | -1.275 | 0.202 |  | R²c: 0.96 |
| Sequencing Run: 2025^b^ | -0.260 | 0.043 | -6.023 | 1.7e-09* |  |  |
| Year: 2023^c^ | 0.180 | 0.042 | 4.299 | 1.7e-05* |  |  |
| Year: 2024^c^ | 0.183 | 0.042 | 4.347 | 1.4e-05* |  |  |

Reference category:  ^a^ LWED; ^b^ 2023; ^c^ 2022

**AICc values for Model Shannon:**

Null model: -32.2

Alternative model: -30.5

**AICc values for Model Richness:**

Null model: 2390.1

Alternative model: 2349.9

**Suppl. Table 3.** Results from LMMs predicting two measures of alpha diversity calculated from VOCs of 134 samples:

- Intensity Model: *Odor Intensity ~ Tamarin species + Year of data collection + Capture day + (1 | Animal identity).*

Data distribution: Gaussian with an identity function

- Shannon Model: *log(Shannon index) ~ Tamarin species + Year of data collection + Capture day + (1 | Animal identity).*

Data distribution: Gaussian with an identity function

Significance: p<0.05 (*), p<0.01 (**), p<0.001 (***). The AICc values for each alternative model and their respective null models are reported at the bottom of the table; for odor intensity, the alternative model outperformed the null model; for Shannon index, the models were equivalent.

| **Intensity Model: Total odor intensity (log-transformed sum of VOC relative peak areas in the sample) as the response variable** | | | | | | |
| --- | --- | --- | --- | --- | --- | --- |
| **Explanatory variable** | **Estimate** | **SE** | ***t-value*** | ***p-value*** | **Random effect: Animal ID** | **R² values** |
| Intercept | 13.654 | 0.273 | 50.009 | < 0.001*** | σ^2^: 0.238  sd: 0.487 | R²m: 0.466 |
| Species: TSUB^a^ | 0.033 | 0.242 | 0.137 | 0.891 |  | R²c: 0.539 |
| Year: 2023^b^ | -2.398 | 0.482 | -4.967 | < 0.001*** |  |  |
| Year: 2024^b^ | -2.381 | 0.893 | -2.667 | 0.009** |  |  |
| Capture day | -0.006 | 0.026 | -0.210 | 0.834 |  |  |
| **Shannon Model: Shannon index on VOC diversity as the response variable** | | | | | | |
| **Explanatory variable** | **Estimate** | **SE** | ***t-value*** | ***p-value*** | **Random effect: Animal ID** | **R² values** |
| Intercept | -0.193 | 0.130 | -1.475 | 0.143 | σ^2^: 0.000  sd: 0.000 | R²m: 0.065 |
| Species: TSUB^a^ | -0.133 | 0.111 | -1.197 | 0.234 |  | R²c: 0.065 |
| Year: 2023^b^ | 0.522 | 0.233 | 2.235 | 0.027* |  |  |
| Year: 2024^b^ | 0.732 | 0.428 | 1.710 | 0.090 |  |  |
| Capture day | -0.015 | 0.013 | -1.195 | 0.234 |  |  |

Reference category:  ^a^ LWED; ^b^ 2022

**AICc values for Model Intensity:**

Null model: 541.1

Alternative model: 463.3

**AICc values for Model Shannon:**

Null model: 269.4

Alternative model: 268.0

**Suppl. Table 4.** Results of PERMANOVA analyses (999 permutations) examining the effects of tamarin species on the Aitchison distances in microbial taxonomic and metabolic pathway compositions, respectively. The models accounted for sequencing run, stratified by year of collection. Significant effects (*p* < 0.05) are indicated with an asterisk.

| **Resp variable** | **Predictors** | **R2** | **F** | **p-value** |
| --- | --- | --- | --- | --- |
| Aitchison distance of microbial taxonomy | Tamarin Species | 0.059 | 8.382 | 0.001* |
|  | Sequencing run | 0.017 | 2.378 | 0.065 |
| Aitchison distance of microbial metabolic pathways | Tamarin Species | 0.010 | 1.351 | 0.178 |
|  | Sequencing run | 0.010 | 1.388 | 0.764 |

**Suppl. Table 5.** Results of RDA analyses (999 permutations) examining the effects of tamarin species on the Aitchison distances in VOC relative abundances. The model accounted for capture day and was stratified by year of collection. Significant effects (p < 0.05) are indicated with an asterisk.

| **Resp. variable** | **Predictors** | **df** | **Variance** | **F** | **p-value** |
| --- | --- | --- | --- | --- | --- |
| Aitchison distance of VOC relative abundance | Tamarin Species | 1 | 0.418 | 0.957 | 0.556 |
|  | Capture # (numeric) | 1 | 1.297 | 2.965 | 0.896 |
|  | Residual | 131 | 57.285 |  |  |

**Suppl. Table 6.** Results from GLMMs predicting two measures of alpha diversity calculated from microbial taxonomy of *Leontocebus weddelli* (n = 76 samples):

- Shannon Model: *Shannon index ~ Group membership + Sequencing year + Year of data collection + Read count per sample + (1 | Animal identity).*

Data distribution: Gamma with a log-link function.

- Richness Model: *Richness ~ Group membership + Sequencing year + Year of data collection + Read count per sample + (1 | Animal identity).*

Data distribution: Poisson with a log-link function.

Significance: p<0.05 (*), p<0.01 (**), p<0.001 (***). The AICc values for each alternative model and their respective null models are reported at the bottom of the table. For the Shannon model, the null model provided a better fit than the alternative model, whereas for the Richness model, the alternative model outperformed the null model.

|  | **Shannon Model: microbial taxonomy** | | | | **Richness Model: microbial taxonomy** | | | |
| --- | --- | --- | --- | --- | --- | --- | --- | --- |
| **GroupID** | **emmean** | **SE** | **LCL** | **UCL** | **emmean** | **SE** | **LCL** | **UCL** |
| AR | 1.940 | 0.049 | 1.840 | 2.030 | 8.879 | 0.008 | 8.863 | 8.895 |
| GD | 1.880 | 0.067 | 1.750 | 2.010 | 8.876 | 0.011 | 8.854 | 8.898 |
| GP | 1.900 | 0.054 | 1.790 | 2.000 | 8.852 | 0.009 | 8.833 | 8.870 |
| IL | 1.920 | 0.048 | 1.830 | 2.010 | 8.880 | 0.008 | 8.865 | 8.896 |
| S7 | 1.980 | 0.064 | 1.850 | 2.100 | 8.877 | 0.011 | 8.855 | 8.899 |
| SS | 1.940 | 0.050 | 1.840 | 2.040 | 8.873 | 0.009 | 8.856 | 8.889 |
| WF | 1.930 | 0.053 | 1.820 | 2.030 | 8.883 | 0.009 | 8.865 | 8.901 |
| **Pair** | **Estimate** | **SE** | **z.ratio** | **p.value** | **Estimate** | **SE** | **z.ratio** | **p.value** |
| AR-GD | 0.003 | 0.014 | 0.247 | 1.000 | 0.003 | 0.014 | 0.247 | 1.000 |
| AR-GP | 0.028 | 0.012 | 2.241 | 0.274 | 0.028 | 0.012 | 2.241 | 0.274 |
| AR-IL | -0.001 | 0.010 | -0.093 | 1.000 | -0.001 | 0.010 | -0.093 | 1.000 |
| AR-S7 | 0.002 | 0.014 | 0.166 | 1.000 | 0.002 | 0.014 | 0.166 | 1.000 |
| AR-SS | 0.007 | 0.012 | 0.564 | 0.998 | 0.007 | 0.012 | 0.564 | 0.998 |
| AR-WF | -0.004 | 0.012 | -0.339 | 0.999 | -0.004 | 0.012 | -0.339 | 0.999 |
| GD-GP | 0.024 | 0.014 | 1.720 | 0.602 | 0.024 | 0.014 | 1.720 | 0.602 |
| GD-IL | -0.004 | 0.012 | -0.345 | 0.999 | -0.004 | 0.012 | -0.345 | 0.999 |
| GD-S7 | -0.001 | 0.015 | -0.074 | 1.000 | -0.001 | 0.015 | -0.074 | 1.000 |
| GD-SS | 0.003 | 0.014 | 0.238 | 1.000 | 0.003 | 0.014 | 0.238 | 1.000 |
| GD-WF | -0.007 | 0.014 | -0.541 | 0.998 | -0.007 | 0.014 | -0.541 | 0.998 |
| GP-IL | -0.028 | 0.012 | -2.414 | 0.193 | -0.028 | 0.012 | -2.414 | 0.193 |
| GP-S7 | -0.025 | 0.014 | -1.791 | 0.554 | -0.025 | 0.014 | -1.791 | 0.554 |
| GP-SS | -0.021 | 0.013 | -1.665 | 0.639 | -0.021 | 0.013 | -1.665 | 0.639 |
| GP-WF | -0.032 | 0.013 | -2.463 | 0.173 | -0.032 | 0.013 | -2.463 | 0.173 |
| IL-S7 | 0.003 | 0.013 | 0.247 | 1.000 | 0.003 | 0.013 | 0.247 | 1.000 |
| IL-SS | 0.008 | 0.011 | 0.667 | 0.994 | 0.008 | 0.011 | 0.667 | 0.994 |
| IL-WF | -0.003 | 0.011 | -0.278 | 1.000 | -0.003 | 0.011 | -0.278 | 1.000 |
| S7-SS | 0.004 | 0.014 | 0.319 | 0.999 | 0.004 | 0.014 | 0.319 | 0.999 |
| S7-WF | -0.006 | 0.014 | -0.454 | 0.999 | -0.006 | 0.014 | -0.454 | 0.999 |
| SS-WF | -0.011 | 0.012 | -0.862 | 0.978 | -0.011 | 0.012 | -0.862 | 0.978 |

**AICc values for Model Shannon:**

Null model: 194.4

Alternative model: 208.0

**AICc values for Model Richness:**

Null model: 1381.8

Alternative model: 1313.9

**Suppl. Table 7.** Results from GLMMs predicting two measures of alpha diversity calculated from microbial taxonomy of *Tamarinus subgrisescens* (n = 59 samples):

- Shannon Model: *Shannon index ~ Group membership + Sequencing year + Year of data collection + Read count per sample + (1 | Animal identity).*

Data distribution: Gamma with a log-link function.

- Richness Model: *Richness ~ Group membership + Sequencing year + Year of data collection + Read count per sample + (1 | Animal identity).*

Data distribution: Poisson with a log-link function.

Significance: p<0.05 (*), p<0.01 (**), p<0.001 (***). The AICc values for each alternative model and their respective null models are reported at the bottom of the table. For the Shannon model, the null model provided a better fit than the alternative model, whereas for the Richness model, the alternative model outperformed the null model.

|  | **Model Shannon: microbial taxonomy** | | | | **Model Richness: microbial taxonomy** | | | |
| --- | --- | --- | --- | --- | --- | --- | --- | --- |
| **GroupID** | **emmean** | **SE** | **LCL** | **UCL** | **emmean** | **SE** | **LCL** | **UCL** |
| IC | 2.00 | 0.029 | 1.94 | 2.06 | 8.875 | 0.004 | 8.868 | 8.883 |
| JI | 1.84 | 0.045 | 1.76 | 1.93 | 8.875 | 0.005 | 8.866 | 8.884 |
| MI | 1.93 | 0.041 | 1.85 | 2.01 | 8.877 | 0.005 | 8.868 | 8.887 |
| OI | 2.00 | 0.030 | 1.94 | 2.06 | 8.859 | 0.005 | 8.851 | 8.868 |
| QB | 1.97 | 0.045 | 1.88 | 2.05 | 8.882 | 0.007 | 8.869 | 8.896 |
| WI | 1.95 | 0.035 | 1.88 | 2.02 | 8.876 | 0.005 | 8.866 | 8.886 |
| YMCM | 1.97 | 0.042 | 1.88 | 2.05 | 8.870 | 0.005 | 8.861 | 8.880 |
| **Pair** | **Estimate** | **SE** | **z.ratio** | **p.value** | **Estimate** | **SE** | **z.ratio** | **p.value** |
| IC-JI | 0.157 | 0.052 | 3.018 | 0.041 | 3.16e-05 | 0.006 | 0.006 | 1.000 |
| IC-MI | 0.071 | 0.047 | 1.535 | 0.724 | -0.002 | 0.006 | -0.380 | 1.000 |
| IC-OI | -0.002 | 0.040 | -0.044 | 1.000 | 0.016 | 0.006 | 2.741 | 0.088 |
| IC-QB | 0.033 | 0.052 | 0.629 | 0.996 | -0.007 | 0.008 | -0.897 | 0.973 |
| IC-WI | 0.051 | 0.043 | 1.188 | 0.899 | -0.001 | 0.006 | -0.111 | 1.000 |
| IC-YMCM | 0.035 | 0.048 | 0.735 | 0.990 | 0.005 | 0.006 | 0.871 | 0.977 |
| JI-MI | -0.086 | 0.057 | -1.500 | 0.745 | -0.002 | 0.006 | -0.357 | 1.000 |
| JI-OI | -0.159 | 0.052 | -3.026 | 0.040 | 0.016 | 0.006 | 2.513 | 0.154 |
| JI-QB | -0.124 | 0.062 | -2.000 | 0.414 | -0.007 | 0.008 | -0.869 | 0.977 |
| JI-WI | -0.106 | 0.054 | -1.975 | 0.431 | -0.001 | 0.006 | -0.111 | 1.000 |
| JI-YMCM | -0.122 | 0.058 | -2.083 | 0.363 | 0.005 | 0.006 | 0.799 | 0.985 |
| MI-OI | -0.073 | 0.046 | -1.589 | 0.690 | 0.018 | 0.006 | 2.804 | 0.075 |
| MI-QB | -0.039 | 0.058 | -0.668 | 0.994 | -0.005 | 0.008 | -0.572 | 0.998 |
| MI-WI | -0.021 | 0.047 | -0.445 | 0.999 | 0.002 | 0.006 | 0.244 | 1.000 |
| MI-YMCM | -0.036 | 0.051 | -0.706 | 0.992 | 0.007 | 0.006 | 1.172 | 0.905 |
| OI-QB | 0.034 | 0.051 | 0.673 | 0.994 | -0.023 | 0.008 | -2.853 | 0.066 |
| OI-WI | 0.052 | 0.033 | 1.588 | 0.690 | -0.017 | 0.007 | -2.488 | 0.164 |
| OI-YMCM | 0.037 | 0.048 | 0.774 | 0.987 | -0.011 | 0.006 | -1.739 | 0.589 |
| QB-WI | 0.018 | 0.055 | 0.322 | 1.000 | 0.006 | 0.008 | 0.750 | 0.989 |
| QB-YMCM | 0.003 | 0.059 | 0.044 | 1.000 | 0.012 | 0.008 | 1.478 | 0.758 |
| WI-YMCM | -0.015 | 0.049 | -0.309 | 1.000 | 0.006 | 0.006 | 0.878 | 0.976 |

**AICc values for Model Shannon:**

Null model: 67.5

Alternative model: 70.5

**AICc values for Model Richness:**

Null model: 751.4

Alternative model: 731.9

**Suppl. Table 8.** Results from GLMMs predicting two measures of alpha diversity calculated from microbial metabolic pathways of *Leontocebus weddelli* (n = 76 samples):

- Shannon Model: *Shannon index ~ Group membership + Sequencing year + Year of data collection + (1 | Animal identity).*

Data distribution: Gamma with a log-link function.

- Richness Model: *Richness ~ Group membership + Sequencing year + Year of data collection + (1 | Animal identity).*

Data distribution: Poisson with a log-link function.

Significance: p<0.05 (*), p<0.01 (**), p<0.001 (***). The AICc values for each alternative model and their respective null models are reported at the bottom of the table. For the Shannon model, the null model provided a better fit than the alternative model, whereas for the Richness model, the alternative model outperformed the null model.

|  | **Model Shannon: metabolic pawhays** | | | | **Model Richness: metabolic pawhays** | | | |
| --- | --- | --- | --- | --- | --- | --- | --- | --- |
| **GroupID** | **emmean** | **SE** | **LCL** | **UCL** | **emmean** | **SE** | **LCL** | **UCL** |
| AR | 1.680 | 0.015 | 1.650 | 1.710 | 5.910 | 0.066 | 5.780 | 6.040 |
| GD | 1.680 | 0.021 | 1.640 | 1.730 | 5.950 | 0.101 | 5.750 | 6.150 |
| GP | 1.680 | 0.017 | 1.650 | 1.720 | 5.920 | 0.089 | 5.750 | 6.100 |
| IL | 1.690 | 0.015 | 1.660 | 1.710 | 5.870 | 0.064 | 5.750 | 6.000 |
| S7 | 1.700 | 0.020 | 1.660 | 1.740 | 5.930 | 0.099 | 5.740 | 6.130 |
| SS | 1.690 | 0.017 | 1.660 | 1.730 | 5.980 | 0.083 | 5.820 | 6.140 |
| WF | 1.680 | 0.017 | 1.650 | 1.710 | 5.840 | 0.085 | 5.670 | 6.000 |
| **Pair** | **Estimate** | **SE** | **z.ratio** | **p.value** | **Estimate** | **SE** | **z.ratio** | **p.value** |
| AR-GD | -0.005 | 0.025 | -0.184 | 1.000 | -0.042 | 0.120 | -0.354 | 1.000 |
| AR-GP | -0.003 | 0.023 | -0.119 | 1.000 | -0.015 | 0.111 | -0.137 | 1.000 |
| AR-IL | -0.006 | 0.018 | -0.320 | 1.000 | 0.033 | 0.062 | 0.526 | 0.998 |
| AR-S7 | -0.017 | 0.024 | -0.696 | 0.993 | -0.027 | 0.119 | -0.227 | 1.000 |
| AR-SS | -0.013 | 0.022 | -0.596 | 0.997 | -0.072 | 0.106 | -0.685 | 0.993 |
| AR-WF | -0.002 | 0.022 | -0.102 | 1.000 | 0.072 | 0.107 | 0.673 | 0.994 |
| GD-GP | 0.002 | 0.026 | 0.074 | 1.000 | 0.027 | 0.132 | 0.205 | 1.000 |
| GD-IL | -0.001 | 0.023 | -0.052 | 1.000 | 0.075 | 0.114 | 0.656 | 0.995 |
| GD-S7 | -0.012 | 0.027 | -0.465 | 0.999 | 0.015 | 0.137 | 0.112 | 1.000 |
| GD-SS | -0.009 | 0.026 | -0.330 | 1.000 | -0.030 | 0.129 | -0.234 | 1.000 |
| GD-WF | 0.002 | 0.025 | 0.096 | 1.000 | 0.114 | 0.128 | 0.888 | 0.974 |
| GP-IL | -0.003 | 0.022 | -0.144 | 1.000 | 0.048 | 0.108 | 0.440 | 1.000 |
| GP-S7 | -0.014 | 0.025 | -0.563 | 0.998 | -0.012 | 0.132 | -0.090 | 1.000 |
| GP-SS | -0.011 | 0.024 | -0.446 | 0.999 | -0.057 | 0.121 | -0.473 | 0.999 |
| GP-WF | 0.0005 | 0.024 | 0.019 | 1.000 | 0.087 | 0.122 | 0.712 | 0.992 |
| IL-S7 | -0.011 | 0.023 | -0.488 | 0.999 | -0.060 | 0.114 | -0.520 | 0.999 |
| IL-SS | -0.007 | 0.021 | -0.349 | 1.000 | -0.105 | 0.103 | -1.015 | 0.951 |
| IL-WF | 0.004 | 0.020 | 0.175 | 1.000 | 0.039 | 0.103 | 0.379 | 1.000 |
| S7-SS | 0.004 | 0.025 | 0.150 | 1.000 | -0.045 | 0.128 | -0.355 | 1.000 |
| S7-WF | 0.015 | 0.025 | 0.591 | 0.997 | 0.099 | 0.128 | 0.768 | 0.988 |
| SS-WF | 0.011 | 0.023 | 0.476 | 0.999 | 0.144 | 0.118 | 1.225 | 0.885 |

**AICc values for Model Shannon:**

Null model: -40.9

Alternative model: -25.7

**AICc values for Model Richness:**

Null model: 1316.7

Alternative model: 1293.6

**Suppl. Table 9.** Results from GLMMs predicting two measures of alpha diversity calculated from microbial metabolic pathways of *Tamarinus subgrisescens* (n = 59 samples):

- Shannon Model: *Shannon index ~ Group membership + Sequencing year + Year of data collection + (1 | Animal identity).*

Data distribution: Gamma with a log-link function.

- Richness Model: *Richness ~ Group membership + Sequencing year + Year of data collection + (1 | Animal identity).*

Data distribution: Poisson with a log-link function.

Significance: p<0.05 (*), p<0.01 (**), p<0.001 (***). The AICc values for each alternative model and their respective null models are reported at the bottom of the table. For the Shannon model, the null model provided a better fit than the alternative model, whereas for the Richness model, the alternative model outperformed the null model.

|  | **Model Shannon: metabolic pathways** | | | | **Model Richness: metabolic pathways** | | | |
| --- | --- | --- | --- | --- | --- | --- | --- | --- |
| **GroupID** | **emmean** | **SE** | **LCL** | **UCL** | **emmean** | **SE** | **LCL** | **UCL** |
| IC | 1.670 | 0.018 | 1.640 | 1.710 | 5.760 | 0.084 | 5.600 | 5.930 |
| JI | 1.690 | 0.024 | 1.650 | 1.740 | 5.900 | 0.115 | 5.670 | 6.120 |
| MI | 1.680 | 0.025 | 1.640 | 1.730 | 5.850 | 0.115 | 5.630 | 6.080 |
| OI | 1.670 | 0.020 | 1.630 | 1.710 | 5.820 | 0.083 | 5.660 | 5.980 |
| QB | 1.700 | 0.029 | 1.640 | 1.760 | 5.860 | 0.130 | 5.600 | 6.110 |
| WI | 1.670 | 0.024 | 1.620 | 1.720 | 5.620 | 0.091 | 5.450 | 5.800 |
| YMCM | 1.660 | 0.024 | 1.610 | 1.710 | 5.840 | 0.113 | 5.610 | 6.060 |
| **Pair** | **Estimate** | **SE** | **z.ratio** | **p.value** | **Estimate** | **SE** | **z.ratio** | **p.value** |
| IC-JI | -0.021 | 0.029 | -0.719 | 0.992 | -0.131 | 0.138 | -0.954 | 0.964 |
| IC-MI | -0.012 | 0.028 | -0.414 | 1.000 | -0.088 | 0.131 | -0.677 | 0.994 |
| IC-OI | 0.003 | 0.026 | 0.107 | 1.000 | -0.055 | 0.111 | -0.493 | 0.999 |
| IC-QB | -0.026 | 0.034 | -0.761 | 0.989 | -0.094 | 0.151 | -0.619 | 0.996 |
| IC-WI | 0.001 | 0.028 | 0.048 | 1.000 | 0.139 | 0.115 | 1.203 | 0.893 |
| IC-YMCM | 0.011 | 0.028 | 0.398 | 1.000 | -0.073 | 0.132 | -0.550 | 0.998 |
| JI-MI | 0.010 | 0.033 | 0.294 | 1.000 | 0.043 | 0.153 | 0.281 | 1.000 |
| JI-OI | 0.024 | 0.030 | 0.788 | 0.986 | 0.077 | 0.136 | 0.562 | 0.998 |
| JI-QB | -0.005 | 0.037 | -0.127 | 1.000 | 0.038 | 0.170 | 0.222 | 1.000 |
| JI-WI | 0.022 | 0.033 | 0.688 | 0.993 | 0.270 | 0.140 | 1.932 | 0.459 |
| JI-YMCM | 0.032 | 0.032 | 0.995 | 0.955 | 0.059 | 0.154 | 0.382 | 1.000 |
| MI-OI | 0.014 | 0.029 | 0.489 | 0.999 | 0.034 | 0.127 | 0.265 | 1.000 |
| MI-QB | -0.014 | 0.037 | -0.387 | 1.000 | -0.005 | 0.165 | -0.031 | 1.000 |
| MI-WI | 0.013 | 0.030 | 0.434 | 1.000 | 0.227 | 0.128 | 1.775 | 0.565 |
| MI-YMCM | 0.023 | 0.030 | 0.756 | 0.989 | 0.016 | 0.141 | 0.112 | 1.000 |
| OI-QB | -0.029 | 0.034 | -0.839 | 0.981 | -0.039 | 0.148 | -0.261 | 1.000 |
| OI-WI | -0.001 | 0.028 | -0.052 | 1.000 | 0.194 | 0.074 | 2.624 | 0.119 |
| OI-YMCM | 0.008 | 0.029 | 0.288 | 1.000 | -0.018 | 0.129 | -0.138 | 1.000 |
| QB-WI | 0.027 | 0.037 | 0.734 | 0.991 | 0.232 | 0.154 | 1.504 | 0.742 |
| QB-YMCM | 0.037 | 0.036 | 1.014 | 0.951 | 0.021 | 0.166 | 0.126 | 1.000 |
| WI-YMCM | 0.010 | 0.031 | 0.321 | 1.000 | -0.211 | 0.132 | -1.607 | 0.678 |

**AICc values for Model Shannon:**

Null model: 9.4

Alternative model: 27.5

**AICc values for Model Richness:**

Null model: 1076.8

Alternative model: 1062.0

**Suppl. Table 10.** Results from LMMs predicting two measures of alpha diversity calculated from VOCs of *Leontocebus weddelli* (n = 72 samples):

- Intensity Model: *Intensity ~ Group membership + Year of data collection + (1 | Animal identity).*

Data distribution: Gaussian with an identity link function.

- Shannon Model: *log(Shannon index) ~ Group membership + Year of data collection + (1 | Animal identity).*

Data distribution: Gaussian with an identity link function.

Significance: p<0.05 (*), p<0.01 (**), p<0.001 (***). The AICc values for each alternative model and their respective null models are reported at the bottom of the table. For the Shannon model, the null model provided a better fit than the alternative model, whereas for the Intensity model, the alternative model outperformed the null model.

|  | **Model Intensity: VOCs** | | | | **Model Shannon: VOCs** | | | |
| --- | --- | --- | --- | --- | --- | --- | --- | --- |
| **GroupID** | **emmean** | **SE** | **LCL** | **UCL** | **emmean** | **SE** | **LCL** | **UCL** |
| AR | 11.200 | 0.439 | 10.300 | 12.100 | -0.198 | 0.213 | -0.631 | 0.235 |
| GD | 11.800 | 0.428 | 10.900 | 12.600 | -0.191 | 0.207 | -0.612 | 0.229 |
| GP | 11.400 | 0.407 | 10.600 | 12.300 | 0.061 | 0.199 | -0.346 | 0.468 |
| IL | 12.600 | 0.382 | 11.800 | 13.400 | -0.005 | 0.180 | -0.367 | 0.356 |
| S7 | 12.300 | 0.602 | 11.100 | 13.500 | -0.224 | 0.282 | -0.787 | 0.339 |
| SS | 12.000 | 0.311 | 11.300 | 12.600 | -0.032 | 0.155 | -0.357 | 0.294 |
| WF | 12.200 | 0.383 | 11.400 | 13.000 | -0.236 | 0.184 | -0.609 | 0.136 |
| **Pair** | **Estimate** | **SE** | **t.ratio** | **p.value** | **Estimate** | **SE** | **z.ratio** | **p.value** |
| AR-GD | -0.519 | 0.602 | -0.862 | 0.976 | -0.007 | 0.292 | -0.023 | 1.000 |
| AR-GP | -0.189 | 0.597 | -0.317 | 1.000 | -0.259 | 0.290 | -0.892 | 0.971 |
| AR-IL | -1.359 | 0.588 | -2.310 | 0.264 | -0.193 | 0.282 | -0.685 | 0.993 |
| AR-S7 | -1.010 | 0.731 | -1.381 | 0.810 | 0.026 | 0.346 | 0.075 | 1.000 |
| AR-SS | -0.721 | 0.536 | -1.346 | 0.823 | -0.166 | 0.262 | -0.634 | 0.995 |
| AR-WF | -0.973 | 0.589 | -1.651 | 0.652 | 0.038 | 0.284 | 0.135 | 1.000 |
| GD-GP | 0.330 | 0.589 | 0.561 | 0.998 | -0.252 | 0.287 | –0.881 | 1.973 |
| GD-IL | -0.840 | 0.580 | -1.448 | 0.773 | -0.186 | 0.278 | -0.671 | 0.994 |
| GD-S7 | -0.490 | 0.725 | -0.676 | 0.993 | 0.033 | 0.343 | 0.096 | 1.000 |
| GD-SS | -0.202 | 0.527 | -0.383 | 1.000 | -0.160 | 0.258 | -0.618 | 0.996 |
| GD-WF | -0.454 | 0.581 | -0.780 | 0.986 | 0.045 | 0.280 | 0.161 | 1.000 |
| GP-IL | -1.170 | 0.559 | -2.094 | 0.376 | 0.066 | 0.269 | 0.247 | 1.000 |
| GP-S7 | -0.820 | 0.719 | -1.141 | 0.912 | 0.285 | 0.341 | 0.835 | 0.980 |
| GP-SS | -0.532 | 0.513 | -1.038 | 0.939 | 0.093 | 0.252 | 0.368 | 1.000 |
| GP-WF | -0.784 | 0.560 | -1.400 | 0.798 | 0.297 | 0.271 | 1.097 | 0.925 |
| IL-S7 | 0.350 | 0.720 | 0.486 | 0.999 | 0.219 | 0.337 | 0.649 | 0.995 |
| IL-SS | 0.638 | 0.494 | 1.292 | 0.851 | 0.027 | 0.239 | 0.111 | 1.000 |
| IL-WF | 0.387 | 0.538 | 0.718 | 0.991 | 0.231 | 0.257 | 0.901 | 0.971 |
| S7-SS | 0.288 | 0.681 | 0.423 | 1.000 | -0.192 | 0.323 | -0.595 | 0.997 |
| S7-WF | 0.037 | 0.720 | 0.051 | 1.000 | 0.012 | 0.339 | 0.036 | 1.000 |
| SS-WF | -0.252 | 0.495 | -0.508 | 0.999 | 0.205 | 0.242 | 0.847 | 0.978 |

**AICc values for Model Intensity:**

Null model: 294.1

Alternative model: 248.9

**AICc values for Model Shannon:**

Null model: 139.9

Alternative model: 139.4

**Suppl. Table 11.** Results from GLMMs predicting two measures of alpha diversity calculated from VOCs of *Tamarinus subgrisescens* (n = 62 samples):

- Intensity Model: *Intensity ~ Group membership + Year of data collection + (1 | Animal identity).*

Data distribution: Gaussian with an identity link function.

- Shannon Model: *log(Shannon index) ~ Group membership + Year of data collection  + (1 | Animal identity).*

Data distribution: Gaussian with an identity link function.

Significance: p<0.05 (*), p<0.01 (**), p<0.001 (***). The AICc values for each alternative model and their respective null models are reported at the bottom of the table. For the Shannon model, the null model provided a better fit than the alternative model, whereas for the Intensity model, the alternative model outperformed the null model.

|  | **Model Intensity: VOCs** | | | | **Model Shannon: VOCs** | | | |
| --- | --- | --- | --- | --- | --- | --- | --- | --- |
| **GroupID** | **emmean** | **SE** | **LCL** | **UCL** | **emmean** | **SE** | **LCL** | **UCL** |
| IC | 11.800 | 0.413 | 10.900 | 12.600 | -0.161 | 0.197 | -0.563 | 0.241 |
| JI | 11.500 | 0.519 | 10.400 | 12.600 | -0.039 | 0.240 | -0.547 | 0.469 |
| MI | 11.700 | 0.485 | 10.800 | 12.700 | -0.287 | 0.234 | -0.770 | 0.195 |
| OI | 12.500 | 0.462 | 11.600 | 13.400 | -0.342 | 0.227 | -0.809 | 0.124 |
| QB | 11.700 | 0.742 | 10.200 | 13.200 | -0.899 | 0.377 | -1.654 | -0.143 |
| WI | 12.100 | 0.530 | 10.000 | 13.100 | -0.280 | 0.256 | -0.800 | 0.240 |
| YM | 12.300 | 0.560 | 10.200 | 13.400 | -0.392 | 0.269 | -0.938 | 0.155 |
| **Pair** | **Estimate** | **SE** | **t.ratio** | **p.value** | **Estimate** | **SE** | **z.ratio** | **p.value** |
| IC-JI | 0.266 | 0.664 | 0.401 | 1.000 | -0.122 | 0.311 | -0.393 | 1.000 |
| IC-MI | 0.027 | 0.634 | 0.043 | 1.000 | 0.126 | 0.305 | 0.416 | 1.000 |
| IC-OI | -0.720 | 0.618 | -1.165 | 0.902 | 0.181 | 0.300 | 0.605 | 0.996 |
| IC-QB | 0.068 | 0.853 | 0.080 | 1.000 | 0.738 | 0.427 | 1.728 | 0.601 |
| IC-WI | -0.281 | 0.669 | -0.421 | 1.000 | 0.119 | 0.322 | 0.371 | 1.000 |
| IC-YM | -0.533 | 0.695 | -0.768 | 0.987 | 0.231 | 0.333 | 0.695 | 0.992 |
| JI-MI | -0.239 | 0.712 | -0.335 | 1.000 | 0.249 | 0.335 | 0.741 | 0.988 |
| JI-OI | -0.986 | 0.695 | -1.419 | 0.787 | 0.303 | 0.331 | 0.917 | 0.966 |
| JI-QB | -0.198 | 0.901 | -0.220 | 1.000 | 0.860 | 0.447 | 1.925 | 0.475 |
| JI-WI | -0.547 | 0.742 | -0.737 | 0.989 | 0.241 | 0.351 | 0.688 | 0.992 |
| JI-YM | -0.799 | 0.764 | -1.046 | 0.938 | 0.353 | 0.360 | 0.981 | 0.954 |
| MI-OI | -0.748 | 0.666 | -1.123 | 0.917 | 0.055 | 0.324 | 0.169 | 1.000 |
| MI-QB | 0.041 | 0.901 | 0.045 | 1.000 | 0.611 | 0.452 | 1.352 | 0.824 |
| MI-WI | -0.309 | 0.710 | -0.435 | 0.999 | -0.007 | 0.343 | -0.021 | 1.000 |
| MI-YM | -0.561 | 0.737 | -0.761 | 0.987 | 0.105 | 0.354 | 0.295 | 1.000 |
| OI-QB | 0.789 | 0.867 | 0.909 | 0.969 | 0.556 | 0.436 | 1.275 | 0.860 |
| OI-WI | 0.439 | 0.687 | 0.639 | 0.995 | -0.062 | 0.340 | -0.182 | 1.000 |
| OI-YM | 0.187 | 0.725 | 0.258 | 1.000 | 0.050 | 0.350 | 0.142 | 1.000 |
| QB-WI | -0.350 | 0.924 | -0.378 | 1.000 | -0.618 | 0.460 | -1.343 | 0.828 |
| QB-YM | -0.602 | 0.939 | -0.641 | 0.995 | -0.507 | 0.466 | -1.087 | 0.929 |
| WI-YM | -0.252 | 0.767 | -0.328 | 1.000 | 0.112 | 0.369 | 0.303 | 1.000 |

**AICc values for Model Intensity:**

Null model: 253.0

Alternative model: 227.6

**AICc values for Model Shannon:**

Null model: 132.9

Alternative model: 143.9

**Suppl. Table 12.** Results of PERMANOVA analyses (999 permutations) examining the effects of group membership on the Aitchison distances in microbial taxonomic and metabolic pathway composition in the two tamarin species separately. The models accounted for sequencing run, stratified by year of collection. Significant effects (*p* < 0.05) are indicated with an asterisk.

| **LWED** | | | | |
| --- | --- | --- | --- | --- |
| **Resp variable** | **Predictors** | **R2** | **F** | **p-value** |
| Aitchison distance of microbial taxonomy | Group ID | 0.124 | 1.670 | 0.009* |
|  | Sequencing run | 0.032 | 2.630 | 0.070 |
| Aitchison distance of microbial metabolic pathways | Group ID | 0.089 | 1.133 | 0.217 |
|  | Sequencing run | 0.015 | 1.177 | 0.153 |
| **TSUB** | | | | |
| **Resp variable** | **Predictors** | **R2** | **F** | **p-value** |
| Aitchison distance of microbial taxonomy | Group ID | 0.113 | 1.135 | 0.239 |
|  | Sequencing run | 0.036 | 2.170 | 0.552 |
| Aitchison distance of microbial metabolic pathways | Group ID | 0.076 | 0.745 | 0.939 |
|  | Sequencing run | 0.054 | 3.158 | 0.106 |

**Suppl. Table 13**. Results of RDA analyses (999 permutations) examining the effects of group membership on the Aitchison distances in VOC relative abundances in the two tamarin species separately. The model was stratified by year of collection. Significant effects (*p* < 0.05) are indicated with an asterisk.

| **Resp. variable** | **Predictors** | **Df** | **Variance** | **F** | **p-value** |
| --- | --- | --- | --- | --- | --- |
| Aitchison distance of VOC relative abundance for **LWED** | Group ID | 6 | 5.070 | 1.018 | 0.515 |
|  | Residual | 65 | 53.930 |  |  |
| Aitchison distance of VOC relative abundance for **TSUB** | Group ID | 6 | 6.200 | 1.165 | 0.036* |
|  | Residual | 55 | 48.800 |  |  |

**Suppl. Table 14.** Classification of mean error rates (%) across resampling iterations of the model performances calculated for the two multiblock-sPLS-DA models linking microbial features (taxonomic or metabolic pathway composition) with VOC profiles. Error rates reported separately for each data block and for each latent component.

| **Multi-Block sPLS-DA microbial taxa - VOCs** | | |
| --- | --- | --- |
|  | Taxa block: Error rate (%) | Odor block: Error rate (%) |
| Component 1 | 3.32% | 47.37% |
| Component 2 | 1.09% | 49.54% |
| **Multi-Block sPLS-DA microbial pathways - VOCs** | | |
|  | Pathways block: Error rate (%) | Odor block: Error rate (%) |
| Component 1 | 25.33% | 44.18% |
| Component 2 | 25.95% | 46.09% |
